# A Time-Sequenced Three-Drug-Loaded Microneedle Patch for the Prevention of Vein Graft Restenosis After Coronary Artery Bypass Grafting

**DOI:** 10.64898/2026.05.20.726718

**Authors:** Bolin Wang, Xiaohang Ding, Longsheng Dai, Yang Yu

**Affiliations:** Capital Medical University Affiliated Anzhen Hospital, Department of Cardiac Surgery, Beijing, China

**Keywords:** coronary artery bypass grafting, microneedle drug delivery system, vein graft restenosis, Transplant vein protection device, PKM2-BCL2

## Abstract

Coronary artery bypass grafting (CABG) is the recommended treatment for severe coronary heart disease; however, restenosis of great saphenous vein grafts continues to limit long-term outcomes because of early thrombosis, mid-term intimal hyperplasia, and late atherosclerosis. To address these sequential pathological processes, this study develops a perivascular drug-loaded microneedle (MN) patch capable of controlled chronological release of tirofiban, rapamycin, and rosuvastatin. The MN patch exhibits uniform conical microneedles (height: 700–800 µm), nanoparticle size of 125 nm, sufficient mechanical strength (1.07 N/needle) for venous intimal penetration, excellent biocompatibility, and no detectable cytotoxicity or hemolysis. In a mouse model of venous graft transplantation, the device significantly reduces early platelet thrombus formation (23.3% to 10%), alleviates mid-term intimal hyperplasia (intimal thickness: 0.84 mm to 0.45 mm at 4 weeks), and improves long-term graft patency with blood flow increasing from 13.1 to 65.8 mL/min at 32 weeks while reducing long-term mortality. Mechanistically, RNA sequencing and functional analyses demonstrate that excessive glycolysis-driven smooth muscle cell proliferation, coupled with abnormal endothelial cell apoptosis, constitutes the core mechanism underlying mid- to long-term restenosis, which the MN device effectively suppresses. This integrated MN patch provides a localised therapeutic strategy for preventing restenosis and improving long-term CABG outcomes.

**Table of Contents:** 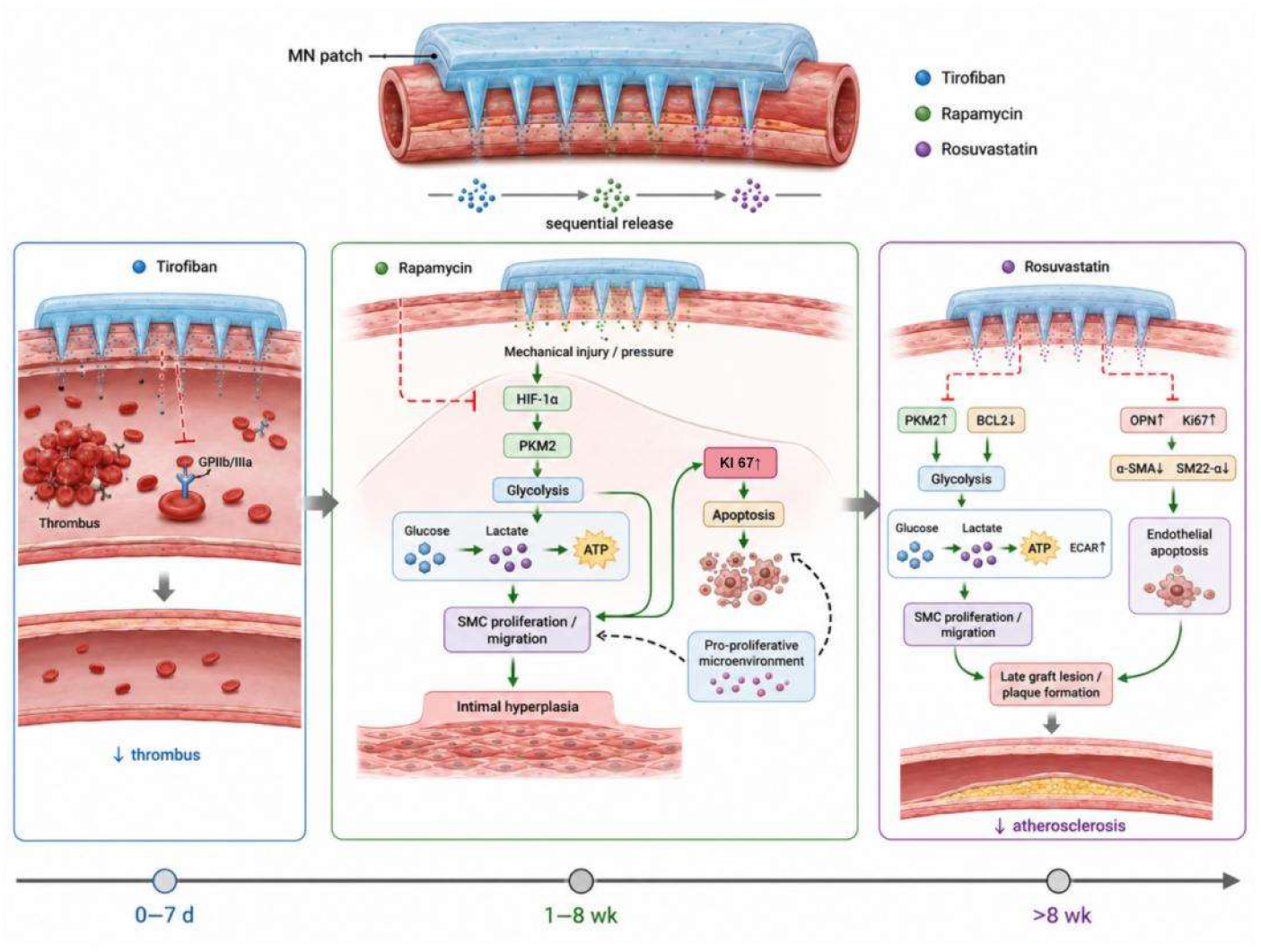

## 1. Introduction

Coronary artery bypass grafting (CABG) is a recommended treatment for severe coronary heart disease (CAD)^[1]^. The patency rate of the great saphenous vein—which is commonly used as a bypass graft in CABG surgery—is approximately 80%,^[2,3]^ and its long-term patency is a major determinant of patient prognosis. However, saphenous vein grafts (SVGs) have a high risk of restenosis,^[4]^ with reported occlusion rates of 3–12% before hospital discharge, failure rates of 8–25% within the first year, and only 50–60% remaining patent after 10 years.^[5–8]^ Advances in surgical techniques and pharmacotherapy have improved mid- and long-term SVG patency rates. Recent studies have reported that composite grafts demonstrate patency comparable to that of arterial grafts at 5 years^[9]^ and SVG patency rates as high as 91% at 8 years.^[10]^ Despite these advances, approximately 13% of patients undergoing CABG require repeat revascularization within 10 years. Furthermore, approximately 18% of all percutaneous coronary interventions (PCIs) are performed in patients with a history of CABG, and approximately 6% of all PCIs are performed on SVGs, highlighting the frequent need for repeat revascularization after CABG.^[11–13]^

SVG stenosis after CABG can be divided into three stages according to its pathophysiological process: During the first week to month after CABG, thrombosis and technical failure are the predominant mechanisms. This is followed by intimal hyperplasia between 1 month and 1 year and atherosclerosis after 1 year.

Early graft failure is attributed to technical factors, such as graft injury during harvesting and anastomotic defects; catheter-related factors, including mismatched catheter size or pre-existing graft pathology; and external factors, such as endothelial cell damage, collagen exposure, platelet adhesion and activation, and exposure of platelet glycoprotein IIb/IIIa (GPIIb/IIIa) receptors, all of which contribute to acute thrombosis. Clinically, ventilator support following CABG often exceeds 10 h, preventing the prompt resumption of oral dual antiplatelet therapy (aspirin combined with clopidogrel or ticagrelor) within the recommended 4–24 h postoperatively.^[14,15]^ Consequently, patients are unable to receive antiplatelet therapy during this period, potentially increasing the risk of early graft failure.

Heparin is commonly used as an anticoagulant to prevent early restenosis of venous grafts following CABG. However, its effects on platelet adhesion, activation, and aggregation are relatively limited,^[16]^ potentially increasing the risk of surgical bleeding. Tirofiban is an injectable GPIIb/IIIa antagonist; however, based on our previous clinical randomized controlled trial,^[17]^ its use increased postoperative bleeding risk in patients undergoing CABG, resulting in poor safety and controllability. To address these clinical challenges, our team developed a controlled-release tirofiban-loaded microneedle system for local intervention in vein grafts during CABG surgery. This system substantially reduced the incidence of perioperative SVG thrombosis after CABG and demonstrated that the proliferative and migratory activities of vein graft vascular smooth muscle cells (VSMCs) are activated during the early postoperative period. This activation may be associated with surgery-induced mechanical injury and the sudden changes in venous pressure.^[18–20]^ Although this drug-loaded microneedle system demonstrated safety and efficacy, its therapeutic effects were limited to thrombosis prevention and treatment. Mid-term restenosis of grafted veins is generally considered to result from intimal hyperplasia,^[21.22]^ which represents an adaptive mechanism known as “arterialization” that occurs within months following CABG. Although this process may lead to mild luminal stenosis, it rarely results in significant early stenosis. Due to the abrupt increase in pressure after graft implantation,^[23–27]^ the vein graft undergoes extensive expansion, resulting in altered traction forces exerted on SMCs and endothelial cells. Simultaneously, activated inflammatory cells secrete a variety of cytokines, including interleukin-1 and interleukin-6, as well as growth factors, such as platelet-derived growth factor and transforming growth factor-beta, which promote the proliferation and migration of VSMCs through phenotypic switching and contribute to excessive proliferation of endothelial cells^[28–32]^.

Although several studies have investigated the molecular biological mechanisms underlying these processes, the specific biomechanical pathways involved remain unclear. The present study further explored the mechanisms associated with lesions occurring during this stage, which initially arise at the anastomotic site and gradually extend throughout the entire SVG over time. Innate immune cells, including mast cells and natural killer cells, also contribute to intimal hyperplasia^[33–36]^. Therefore, intimal hyperplasia can be classified into two major components. One component results from changes in cellular traction forces involving SMCs and endothelial cells caused by excessive vein graft expansion, whereas the other component arises from inflammatory mediators secreted by activated cells during the early postoperative period.

To address excessive vein graft expansion, our team independently developed a hydrogel-based external stent with favorable mechanical properties and ease of application^[37]^. Although this device improved venous graft patency during the first few postoperative months, its efficacy remained suboptimal. Therefore, we focused on a second intervention target for intimal hyperplasia. We subsequently prepared a rapamycin-loaded hydrogel patch. Although incorporation of the drug affected the mechanical properties of the patch^[20]^, the system enabled sustained release of the mammalian target of rapamycin (mTOR) inhibitor rapamycin. The drug-loaded patch significantly improved vein graft patency during the first few postoperative months, and our findings demonstrated that hypoxia inducible factor 1 alpha (HIF-1α) serves as a potential therapeutic target for restenosis following vein graft surgery at the cellular level. However, validation from a cellular biomechanics perspective remains lacking^[20]^.

Intimal hyperplasia of the grafted vein forms the pathological basis of atherosclerosis and may lead to late SVG failure. The progression rate of SVG atherosclerosis is higher than that of primary coronary atherosclerosis. SVG atherosclerosis is generally concentric and diffuse, with a less distinct fibrous cap, making plaques more susceptible to rupture.^[38–44]^ Atherosclerotic changes can be observed 1 year after CABG. The earliest manifestations include foam cell accumulation, followed by the development of necrotic cores, which are typically observed 2–5 years after surgery and often contribute to the formation of intermediate SVG lesions. Subsequently, the necrotic core may cause bleeding and spread within the plaque due to neovascular leakage, ultimately leading to plaque rupture, thrombosis, and SVG occlusion. And the latest RCT findings of the year indicate that the use of PCSK9 inhibitors does not improve long-term patency after CABG surgery^[45]^.Although oxidative stress has been implicated in atherosclerosis, whether SVG atherosclerosis follows mechanisms similar to those of native coronary artery disease, as well as the specific oxidative stress-related molecular pathways involved, remains unclear.

Based on the pathological progression of vein graft restenosis after CABG and our previous findings^[17–20]^, we aimed to develop a comprehensive protective system targeting the entire pathological process of vein graft restenosis. Immediately after surgery, an external wrapping system is applied to limit excessive graft expansion. Early controlled release of tirofiban is used to inhibit perioperative thrombosis, whereas sustained release of rapamycin beginning 1 week after surgery targets medium- and long-term restenosis caused by intimal hyperplasia. In addition, a hydrogel microneedle system loaded with rosuvastatin is designed to provide controlled release beginning 1 month after surgery to inhibit atherosclerosis. This study aims to describe the preparation process of this protective system and evaluate its effects on vein graft restenosis across different pathological stages.

According to distinct mechanisms operating at different stages, the primary endpoint for early restenosis was thrombus incidence, whereas mortality served as the endpoint of mid- to long-term restenosis. In the absence of mortality events, the hemodynamic index—which is directly associated with restenosis and mortality—was used as the surrogate endpoint.

## 2. Results

### 2.1 Physical Properties of the Drug-Loaded Microneedle Device

The demolded microneedle patch array exhibited a square shape, with a side length of 1.5 cm, a flat base, uniform microneedle length, well-defined morphology, and no obvious fractures, indicating favorable physical and mechanical properties of the microneedle patch. As shown in Figures S1B and S1C, the drug-loaded microneedle device consisted of a pyramid-shaped single microneedle with an average height of approximately 700–800 µm and an average width of about 300 µm at the bottom edge, which is consistent with the dimensions of the polydimethylsiloxane (PDMS) template. Figure S1D shows the microstructure of the drug-loaded nanoparticles observed using scanning electron microscope (SEM). The nanoparticles exhibited a uniform spherical morphology with an average diameter of 125 nm. These findings indicate that nanocrystals with uniform particle size were successfully prepared, providing a basis for ensuring the homogeneous drug dispersion within the hydrogel matrix and achieving sustained, controlled, and stable drug release during treatment, as shown in Figure S1E.

Compared with the undoped hyaluronic acid microneedle patch, the tirofiban-loaded microneedle device exhibited distinct infrared absorption peaks at 1,725 and 1,834 cm^-1^, corresponding to the chemical structures of -SO_2_ and -CH_3_ in tirofiban, respectively. The rapamycin-loaded microneedle device exhibited significant characteristic infrared absorption peaks at 2,792 and 2,931 cm^-1^, corresponding to the chemical groups -C=O and -CH_2_-CH_2_-, respectively. These results indicate the successful incorporation of rapamycin into the microneedle device. Rosuvastatin exhibited a significant characteristic infrared absorption peak at 3,961 cm^-1^, corresponding to the chemical group -CaH_2_. Collectively, the infrared spectroscopy results confirmed the successful incorporation of all three drugs into the microneedle device.

Liquid chromatography–tandem mass spectrometry (LC–MS/MS) was used to continuously measure the concentrations of the three drugs in the homogenized transplanted blood vessel samples obtained from animals in the drug-loaded microneedle experimental group. The drug release curves over time are shown in Figure S1F. Tirofiban, which served as the early-stage therapeutic agent, exhibited relatively rapid release, with approximately 37% released within 3 days, 65% within 7 days, and 88% within 2 weeks. Rapamycin release began on day 12, with approximately 12% released on day 14, 54% by week 4, and approximately 85% by week 7. Rosuvastatin release began on day 40, reaching a release rate of 58% by week 20 and 91% by week 35. These results indicate that the drugs could be continuously and gradually released from the microneedle patches, thereby exerting sustained therapeutic effects. As the microneedle tips dissolved, the drugs were completely released, whereas the remaining drug contained within the vascular wall patch gradually diffused and spread to the vascular endothelium.

Mechanical properties were evaluated using an electronic force-testing machine. As shown in Figure S1G, the microneedles exhibited a mechanical strength of 1.07 N per needle. According to previous studies^[18–20]^, this level of force is sufficient to penetrate the vascular intima without causing structural damage to the microneedle device.

Figure S1H shows the immunofluorescence results obtained from experimental animals at 7 days, 4 weeks, and 16 weeks after surgery. At 7 days postoperatively, tirofiban was the predominant released drug, with a small amount of rapamycin also present. At 4 weeks, rapamycin and rosuvastatin represented the primary released drugs. By 16 weeks, rosuvastatin had become the dominant drug, although a small amount of rapamycin remained detectable. These findings indicate that drug distribution within the vascular endothelium became increasingly extensive over time, confirming that microneedle patches loaded with drug nanoparticles effectively penetrated the endothelial layer.

The cytotoxicity of the drug-loaded microneedle devices toward human umbilical vein smooth muscle cells (HUVSMCs) was evaluated after incubation at 37 °C for 72h. As shown in Figure 1I, microneedles loaded with high concentrations of tirofiban did not exhibit significant cytotoxicity. At a drug loading concentration of 50 ug/mL, the lowest toxic inhibitory effect on cellular viability was observed, confirming the favorable cell compatibility of the drug-loaded microneedle device.

**Figure 1.**
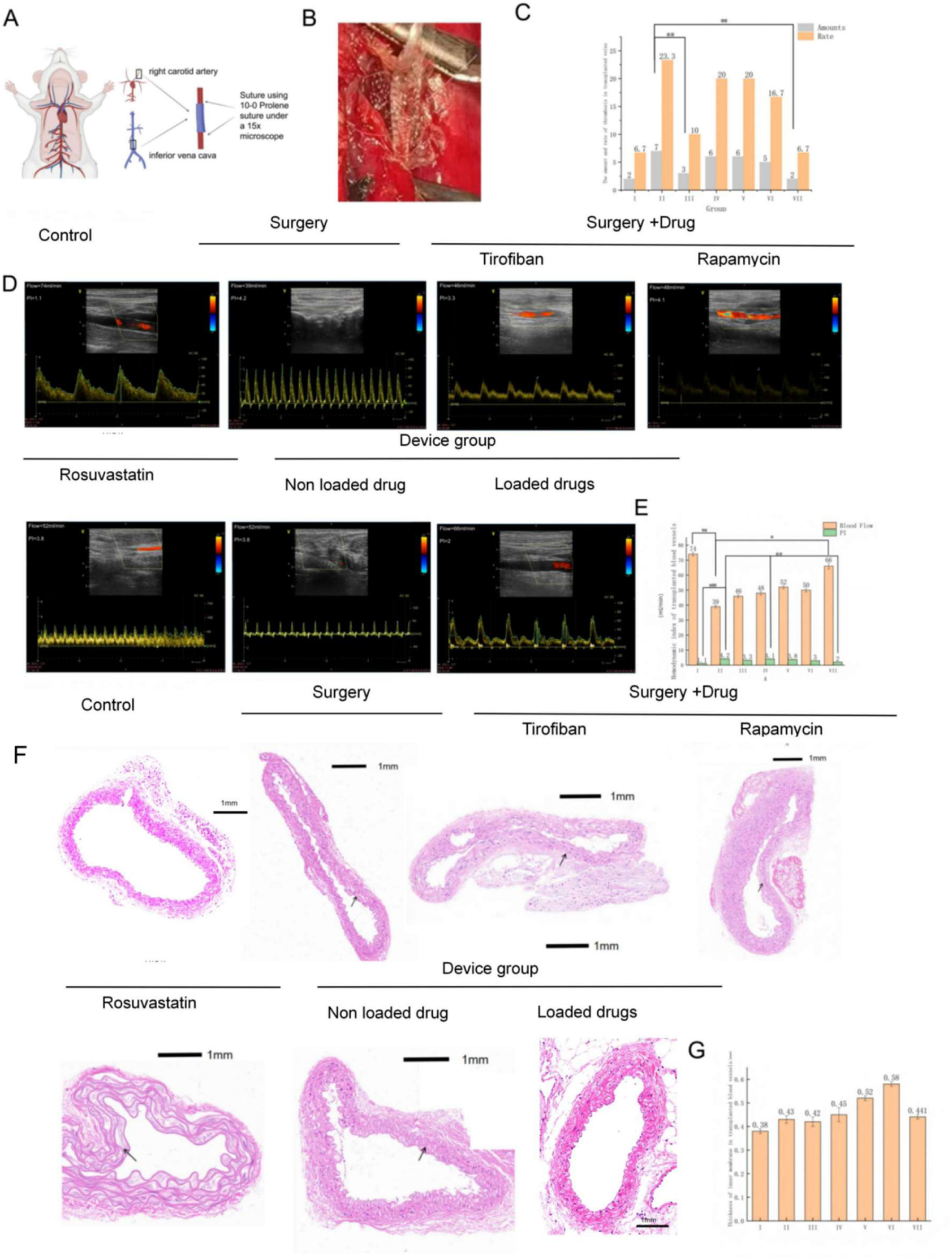
(A) Establishment of a mouse cuff-based vein graft model. (B) Schematic illustration of drug-loaded microneedle device application. (C) Number and incidence of platelet thrombosis in experimental animals. (D–E) Hemodynamic parameters and statistical analyses of experimental animals in each group. (F–G) Representative HE staining images and quantitative analysis of transplanted veins in each group.

The biocompatibility and safety of the hydrogel and drug-loaded hydrogel solutions were evaluated using human blood samples. Both solutions were prepared as microneedle devices and subsequently dissolved prior to testing. The hemolysis rates of the negative control group, hydrogel solution group, drug-loaded hydrogel solution group, and positive control groups were 1.9 ± 0.11, 2.21 ± 0.31, 2.24 ± 0.33, and 99.2 ± 0.54%, respectively. As shown in Figure S1J, the drug-loaded microneedle formulations did not cause hemolysis in human blood samples.

### 2.2 Effect of the Drug-Loaded Microneedle Device on Early Restenosis (Perioperative Thrombosis) in Transplanted Veins

The establishment of the animal model is shown in Figure 1A, and the grouping strategy and model establishment are described in detail in the Methods section. The application of the drug-loaded microneedle device is shown in Figure 1B. The device ompletely surrounded the transplanted blood vessels and enabled drug delivery directly to the vascular endothelium. The number and incidence of platelet thrombosis in each group within 72 h are shown in Figure 1C. Platelet thrombosis occurred in all animal groups. Interestingly, although Group I underwent only cervical incision and closure procedures, two animals developed platelet thrombosis following carotid artery dissection, corresponding to an incidence rate of 6.7%. This observation may be associated with the atherosclerotic characteristics of the experimental animal model. In Group II, platelet thrombus formation occurred in seven animals, corresponding to an incidence of 23.3%, whereas only three animals in Group III developed platelet thrombus, corresponding to an incidence of 10%. Compared with Group II, this difference was statistically significant, indicating that peripheral administration of tirofiban can effectively reduce platelet thrombus formation. However, our previous randomized controlled clinical study in humans demonstrated that systemic tirofiban administration was associated with an increased risk of major bleeding.^[17]^

The number of animals with platelet thrombosis in Groups IV, V, and VI was 6, 6, and 5, respectively, corresponding to improvement rates of 20%, 20%, and 16.7%, respectively. No statistically significant differences were observed compared with the other groups. By contrast, in Group VII—which received the drug-loaded microneedle device—only two animals developed platelet thrombosis, similar to the control group. These results confirm the effectiveness of the drug-loaded microneedle device from a macroscopic perspective.

To further validate these findings, ultrasound examinations were performed before sacrifice to evaluate the hemodynamic parameters of the transplanted blood vessels in the experimental animals. Representative ultrasound images from each group are shown in Figure 1D, and the statistical results are illustrated in Figure E. The blood flow rates of the transplanted blood vessels in Groups I–VII were 74, 39, 46, 48, 52, 50, and 66 mL/min, respectively. Statistically significant differences were observed between Groups I and II, as well as between Groups VII and II. The pulsatility index (PI) values in Groups I–VII were 1.1, 4.2, 3.3, 4.1, 3.8, 3.0, and 2.0, respectively.

Statistically significant differences were observed between Groups I and II and between Groups II and IV. Additionally, statistically significant differences were identified among Groups VII, II, and IV. These findings indicate that the drug-loaded microneedle device improved the early hemodynamic characteristics of transplanted veins. Although the device did not fully restore hemodynamic function to levels comparable with those observed following transplantation surgery alone, the observed improvements were encouraging.

Interestingly, despite the reduction in platelet thrombosis incidence, Group III did not show a corresponding improvement in hemodynamic parameters. This discrepancy may be related to the early release of rapamycin from the microneedle drug delivery device, which may inhibit SMC proliferation and migration. Early SMC proliferation and migration in transplanted veins have been previously reported in our published studies^[18–20]^.

Subsequently, hematoxylin and eosin (HE) staining of transplanted veins was performed to assess intimal compression by assessing intimal thickness. Representative HE staining images are shown in Figure 1F, and quantitative measurements of intimal thickness are shown in Figure 1G. The intimal thicknesses of Groups I–VII were 0.38, 0.43, 0.42, 0.45, 0.52, 0.58, and 0.411 mm, respectively, with no statistically significant differences among the groups. These findings indicate that, although SMC activation occurred during the early postoperative stage, it was insufficient to induce significant intimal thickening. Therefore, early protection of graft veins should primarily focus on preventing platelet thrombosis.

### 2.3 Molecular Mechanisms of the Drug-Loaded Microneedle Device in Early Restenosis (Perioperative Thrombosis) of Transplanted Veins

As shown in Figure 2A and presented in Table 1, the platelet levels in Groups II, IV, V, and VI were consistently higher than those in Groups I, III, and VII. Approximately 72 h after surgery, platelet counts reached near-peak levels in all groups, which may explain why the early postoperative period (72 h after surgery) represents a high-risk stage for perioperative myocardial infarction. The platelet levels in Groups III and V were similar to those in Group 1, indicating that tirofiban and the drug-loaded microneedle devices effectively inhibited platelet thrombus formation.

**Figure 2.**
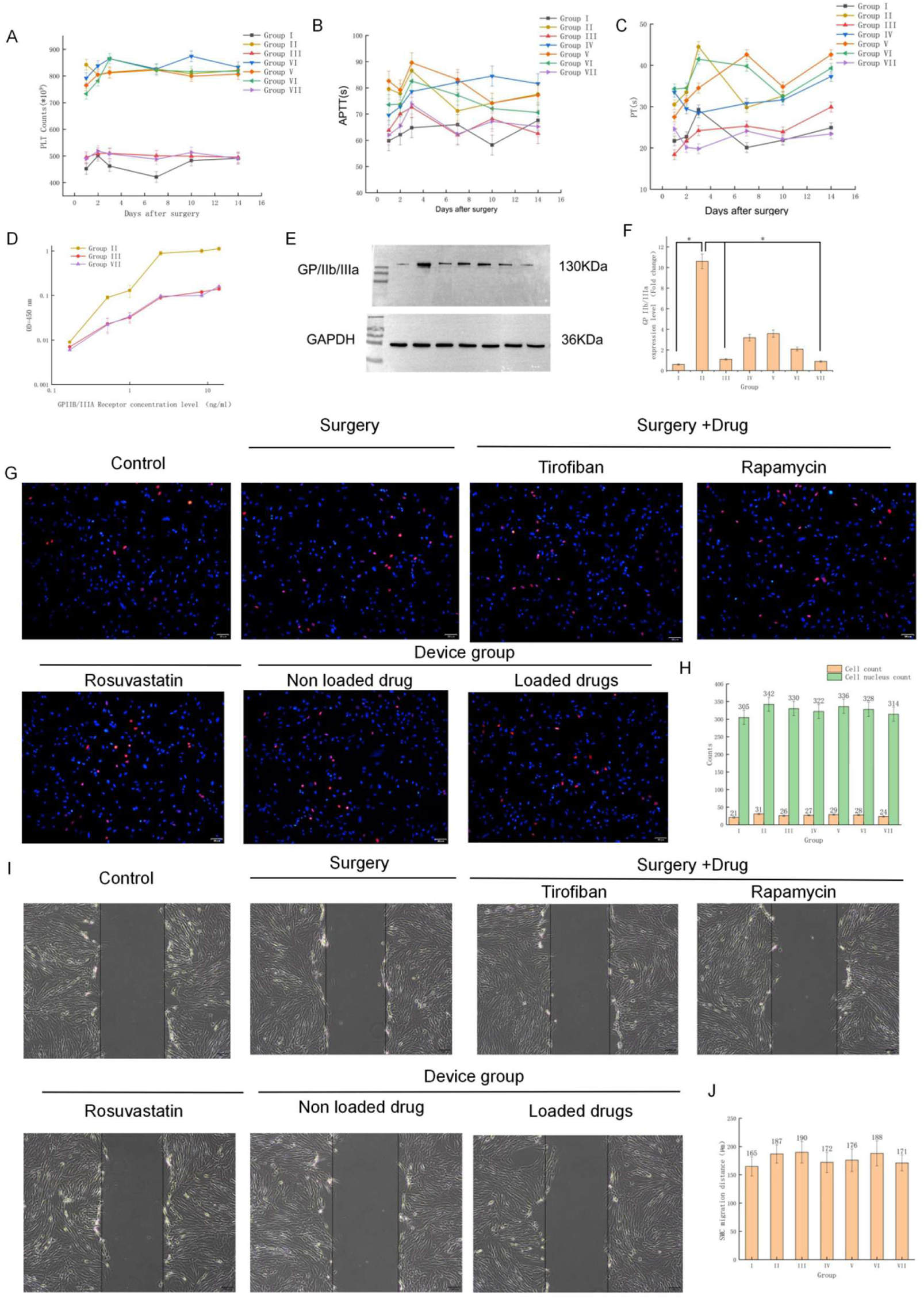
Detection of coagulation-related mechanisms in experimental animals. (A) Platelet levels in each animal group. (B) APTT levels in each group. (C) PT levels in each group. (D) Enzyme-linked immunosorbent assay–based evaluation of GPIIb/IIIa levels in Groups II, III, and VII. (E) Western blot analysis of GPIIb/IIIa expression. (G–H) EdU assay results and quantitative analysis of SMC proliferation in each group. (I–J) Scratch assay results and quantitative analysis of SMC migration in each group.

According to the results shown in Figures 2B and 2C and presented in Table 2, the activated partial thromboplastin time (APTT) and prothrombin time (PT) exhibited trends similar to those observed for PLT levels. These findings suggest that surgery affected the systemic coagulation system of the experimental animals and altered coagulation times. However, following intervention with tirofiban and the drug-loaded microneedle device, APTT and PT at certain postoperative time points were even lower than those observed in the nonsurgical group. These findings support the effectiveness of the drug-loaded microneedle device in regulating platelet activity and coagulation-related outcomes.

As mentioned above, GPIIb/IIIa receptors play a critical role in platelet thrombus formation, whereas tirofiban functions as a GPIIb/IIIa receptor antagonist. Therefore, GPIIb/IIIa expression was evaluated by measuring the optical density (OD) of the drug-loaded microneedle device at a wavelength of 450 ± 2 nm, as shown in Figure 2D. The results showed that when the OD value was 450, Groups III and VII exhibited lower GPIIb/IIIa levels than Group II at an OD of 450 nm. These findings suggest that the interventions administered in Groups III and VII effectively inhibited GP IIb/IIIa activity.

Furthermore, we measured GP IIb/IIIa protein expression levels in each group 72 h after surgery using Western blot analysis. The results are shown in Figures 2E and 3F. GP IIb/IIIa protein expression was significantly increased in Group II, whereas expression levels in Groups I, III, and VII were significantly lower than those observed in Group II. Collectively, these findings experimentally confirm the effective targeting of GP IIb/IIIa receptors by the drug-loaded microneedle device. The proliferation activity of HUVSMCs 72 h after transplantation was evaluated by an 5-ethynyl-2′-deoxyuridine (EdU) assay, and the results are shown in Figures 2G and 2H. Although the numbers of SMCs and cell nuclei in Group II appeared greater than those in the other groups, the differences were not statistically significant. The scratch test results are shown in Figures 2I and 2J. Although Groups I, IV, and VII exhibited relatively lower migration levels than the other groups, these differences did not reach statistical significance.

**Figure 3.**
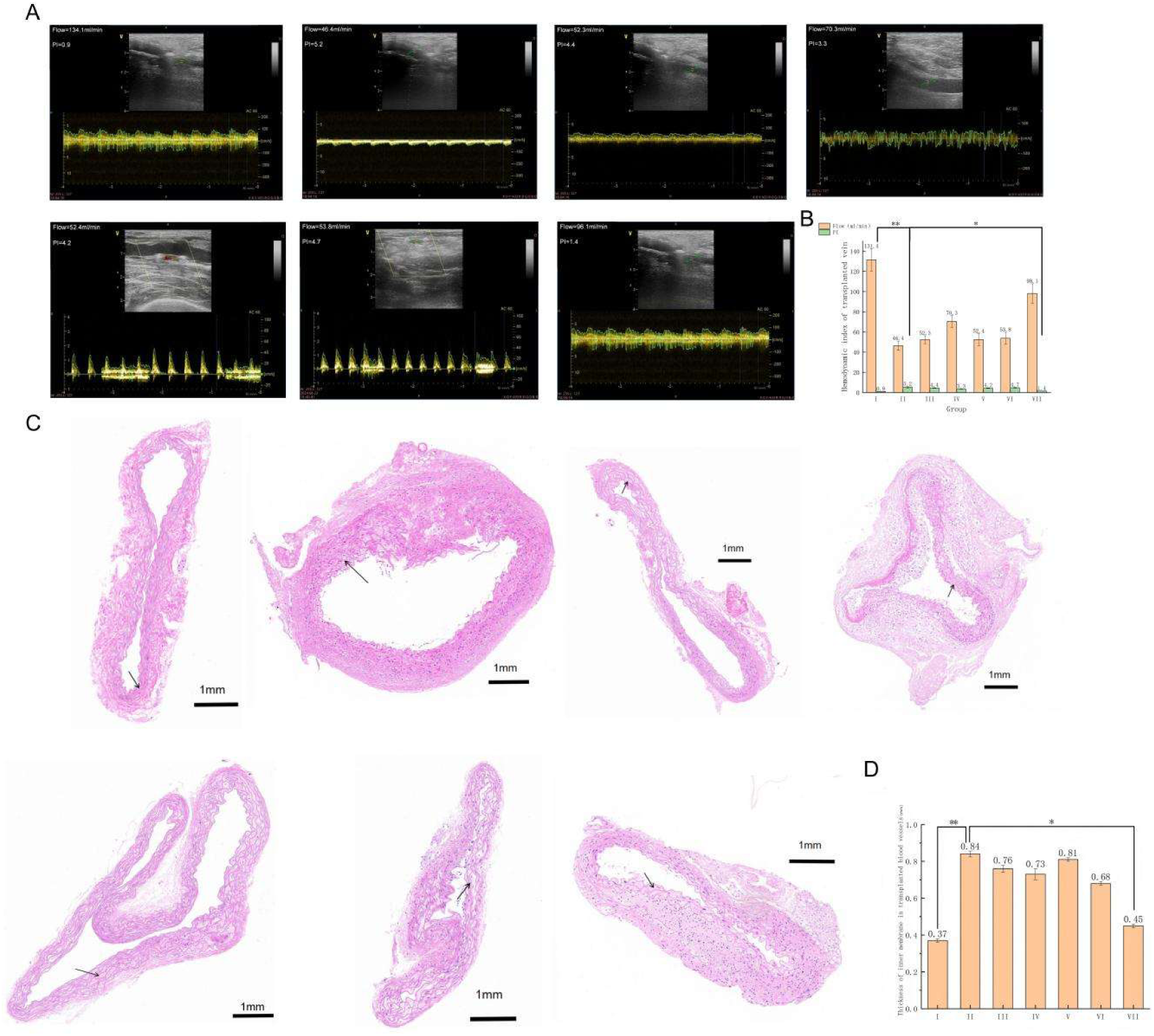
(A–B) Hemodynamic parameters and corresponding statistical analyses of experimental animals at 28 days after surgery. (C–D) Representative HE staining images and quantitative analysis of transplanted veins in experimental animals at 28 days after surgery.

These findings indicate that early restenosis may not be related to the proliferation and migration of VSMCs, although cellular activation appears to occur during the early stage. Rapamycin and the drug-loaded microneedle device effectively inhibited this activation. The EdU and scratch assay results of Group III were higher than those observed in the other groups, further indicating that tirofiban does not exert a protective effect against early restenosis at the cellular level.

| Group | 1 day after surgery | 2 days after surgery | 3 days after surgery | 7 days after Surgery | 10 days after surgery | 14 days after surgery |
| --- | --- | --- | --- | --- | --- | --- |
| I | 432±6.8 | 493±8.3 | 462±7.3 | 433±9.5 | 472±12.9 | 489±12.5 |
| II | 865±17.6 | 796±15.3 | 794±19.3 | 785 ± 16.2 | 779±16.6 | 798±12.6 |
| III | 488±7.8 | 516±11.4 | 508±6.3 | 495±9.2 | 479±8.6 | 491±9.8 |
| IV | 778±13.6 | 822±19.6 | 869±19.3 | 806±13.3 | 871±13.1 | 811±9.9 |
| V | 736±13.6 | 785±8.3 | 802±16.5 | 786±12.5 | 794±16.1 | 796±5.9 |
| VI | 726±8.2 | 784±11.3 | 862±13.4 | 811±10.8 | 804±18.2 | 808±12.2 |
| VII | 476±9.8 | 521±8.6 | 503±6.3 | 486±9.5 | 499±6.8 | 492±10.0 |

| Gro<br>up | 1 day after surgery |  | 2 days after<br>surgery |  | 3 daysafter surgery |  | 7 days after<br>surgery |  | 10 days after<br>surgery |  | 14 days after<br>surgery |  |
| --- | --- | --- | --- | --- | --- | --- | --- | --- | --- | --- | --- | --- |
|  | PT | APTT | PT | APTT | PT | APTT | PT | APTT | PT | APTT | PT | APTT |
| I | 21.4±1.8 | 59.4±6.6 | 22.6±2.6 | 61.8±4.6 | 28.8±2.6 | 62.7±9.3 | 20.4±1.7 | 63.1±5.4 | 19.2±1.7 | 56.3±6.7 | 23.1±2.6 | 64.2±5.6 |
| II | 31.3±2.4 | 79.8±0.9 | 33.2±3.3 | 78.8±9.2 | 46.7±3.5 | 85.4±7.8 | 29.8±3.3 | 71.2±9.3 | 29.2±2.3 | 74.4±9.8 | 38.2±3.3 | 75.4±9.2 |
| III | 17.2±1.4 | 63.4±5.8 | 21.2±1.8 | 69.4±5.6 | 22.5±2.1 | 72.1±6.8 | 23.6±3.6 | 60.9±6.2 | 22.6±2.4 | 63.4±5.6 | 29.1±2.3 | 61.0±5.5 |
| IV | 33.5±3.3 | 69.8±7.1 | 29.2±4.9 | 73.2±8.3 | 31.6±2.3 | 76.9±6.4 | 29.6±3.5 | 80.8±5.3 | 29.8±3.6 | 83.5±6.7 | 34.3±3.1 | 81.9±7.4 |
| V | 29.8±2.1 | 83.6±9.1 | 31.2±2.6 | 78.2±7.7 | 33.6±3.5 | 89.2±9.3 | 44.3±6.2 | 81.3±6.9 | 33.4±3.2 | 76.1±6.9 | 43.2±3.4 | 75.6±4.6 |
| VI | 34.2±4.2 | 74.5±6.4 | 32.8±4.4 | 74.8±6.3 | 43.5±6.3 | 83.4±8.8 | 39.6±3.9 | 76.3±4.8 | 32.5±2.8 | 74.6±8.7 | 38.4±2.9 | 72.6±5.4 |
| VII | 24.2±3.6 | 61.3±5.2 | 19.6±2.2 | 63.8±7.2 | 18.4±2.3 | 73.0±6.6 | 18.8±2.6 | 60.7±5.5 | 19.4±1.6 | 62.8±3.6 | 19.7±1.6 | 62.7±4.0 |

### 2.4 Effect of the Drug-Loaded Microneedle Device on Mid-Term Restenosis of Transplanted Veins

None of the animals died 4 weeks after surgery. To evaluate mid-term patency of the transplanted veins, ultrasound was used to assess the hemodynamic parameters of the transplanted blood vessels in experimental animals. Representative ultrasound results for each group are shown in Figure 3A, and the statistical analyses are shown in Figure 3B. The blood flow rates of transplanted blood vessels from Group I–VII were 131.4, 46.4, 52.3, 70.3, 52.4, 53.8, and 98.1 mL/min, respectively. Statistically significant differences were observed between Groups I and II and between Groups VII and II. Although Group IV exhibited an increase in blood flow compared with Group II, this difference was not statistically significant.

The PI values for groups I–VII were 0.9, 5.2, 4.4, 3.3, 4.2, 4.7, and 1.4, respectively. The statistical significance of these differences was similar to that observed for blood flow measurements. These findings indicate that peripheral administration of rapamycin can improve the hemodynamic characteristics of transplanted veins, although the improvement was not statistically significant. By contrast, the drug-loaded microneedle device significantly improved the mid-term hemodynamic parameters of transplanted veins, further demonstrating the advantages of sustained and targeted drug delivery provided by this device.

Next, we evaluated the intimal thickness of the transplanted veins through HE staining to assess the degree of lumen compression. Representative HE staining images are shown in Figure 3C, and quantitative measurements of intimal thickness are shown in Figure 3D. The intimal thicknesses in Groups I–VII were 0.37, 0.84, 0.76, 0.73, 0.81, 0.68, and 0.45 mm, respectively. Compared with Groups I and VII, group II exhibited significantly greater intimal thickness, and the difference was statistically significant. These findings are consistent with the observed hemodynamic parameters and indicate that reducing SMC proliferation and migration is a key mechanism for suppressing mid-term restenosis. The drug-loaded microneedle protective device achieved substantial therapeutic effects through targeted and sustained drug release.

### 2.5 Molecular Mechanisms of the Drug-Loaded Microneedle Device in Mid-Term Restenosis Intervention of Transplanted Veins

On postoperative day 28, the VSMCs were isolated from the transplanted veins in each experimental group. VSMC proliferation and migration in each group were evaluated using EdU and scratch tests, respectively. The EdU results are shown in Figures 4A and 4B. Both the number of nuclei and proliferating cells were significantly higher in Group II than in Groups I and VII, and the difference was statistically significant. The scratch test results are shown in Figures 4C and 4D and demonstrated trends similar to those observed in the EdU assay. The migration level of VMCs was significantly higher in Group II than in Groups I and VII, and the difference was statistically significant.

**Figure 4.**
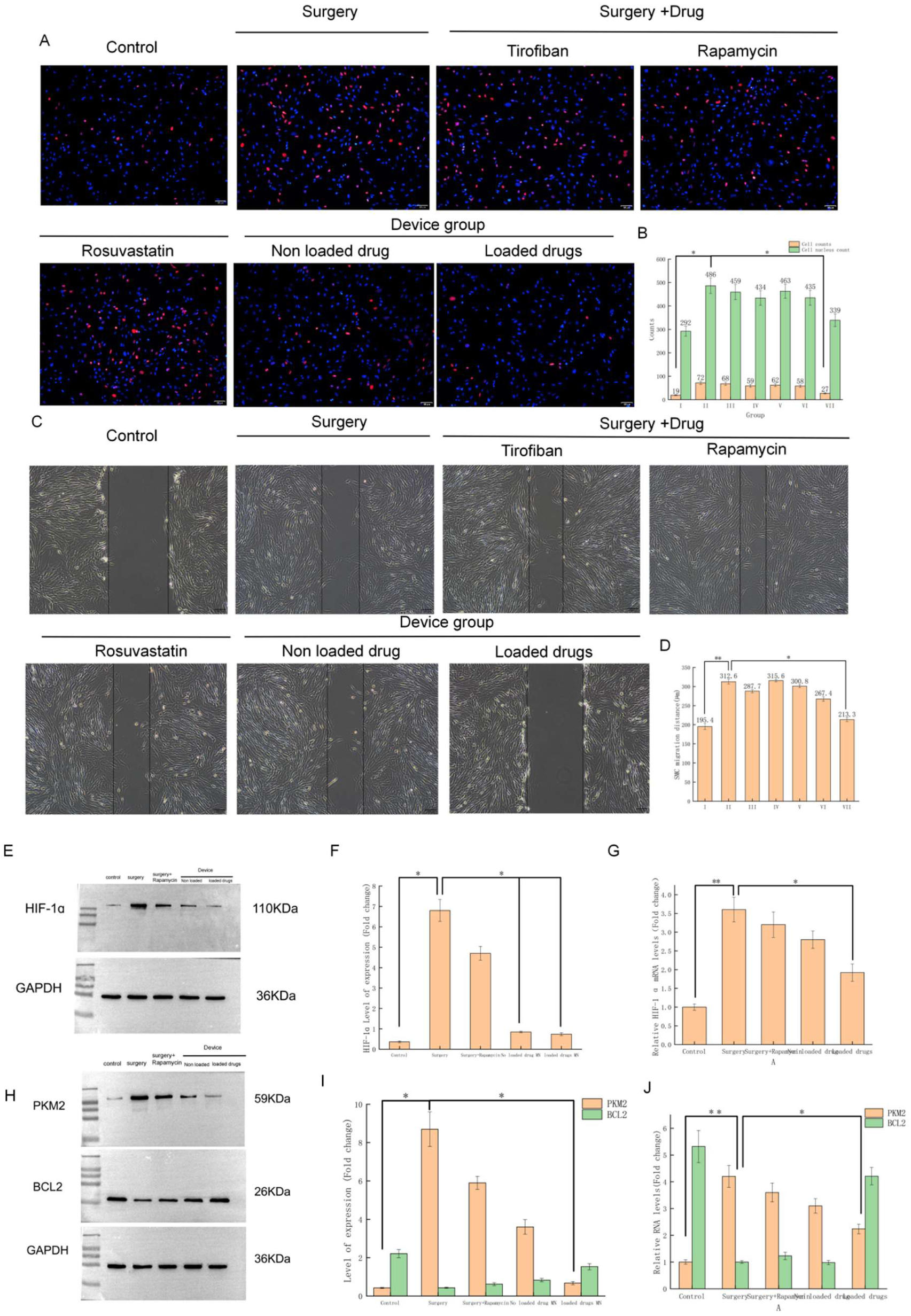
(A–B) EdU assay results and quantitative analysis of SMCs isolated from transplanted veins at 28 days after surgery. (C–D) Scratch test results and quantitative analysis of SMCs extracted from transplanted veins at 28 days after surgery. (E–F) Western blotting results and protein expression analysis of HIF-1α in transplanted veins from each group. (G) qPCR results and mRNA expression analysis of HIF-1α.

Our previous research has shown that the transcription factor HIF-1α is closely associated with mid-term restenosis^[18–20]^. Therefore, we performed Western blotting and quantitative polymerase chain reaction (qPCR) to determine the protein and mRNA expression levels of HIF-1α in Groups I, II, IV, VI, and VII, which were associated with mid-term restenosis. As shown in Figures 4E–G, both the protein and mRNA expression levels of HIF-1α in Group II were significantly higher than those in Groups I and VII, with statistically significant differences.

Previous studies have shown that pyruvate kinase M2 (PKM2) and B-cell lymphoma-2 (BCL2) are associated with long-term restenosis.^[20,24]^ Therefore, we hypothesized that these molecules may also become activated during the mid-term stage. Western blotting and qPCR analyses were used to detect protein and mRNA expression levels of PKM2 and BCL2 in Groups I, II, IV, VI, and VII. As shown in Figures 4H–J, the expression pattern of PKM2 was similar to that of HIF-1α, with significantly increased protein and mRNA levels in Group II compared with Groups I and VII. By contrast, the protein and mRNA levels of BCL2 were significantly lower in Group II than in Groups I and VII. These findings indicate that the molecular factors associated with long-term restenosis may already be activated during the mid-term stage. Furthermore, the results also suggest that the drug-loaded microneedle device effectively inhibits SMC proliferation and migration, thereby suppressing mid-term restenosis of transplanted veins and highlighting the importance of controlled drug release from the device.

Interestingly, a significant difference in HIF-1α protein expression was observed between Groups VI and II. This finding may be related to the intrinsic effects of the microneedle device itself, including its ability to mechanically limit excessive dilation of transplanted veins. Therefore, subsequent biomechanical studies were conducted.

To further investigate the molecular biological mechanisms associated with mid-term restenosis, we performed RNA sequencing on vein graft samples obtained from experimental animals in the control, surgery, and drug-loaded microneedle device groups, with three animals included in each group (Figure5 A). A total of 154 differentially expressed genes (DEGs) were identified between the control and surgery groups, as shown in the volcano plot in Figure5 B. The Kyoto Encyclopedia of Genes and Genomes (KEGG) pathway enrichment results are shown in Figures 5 C and 5D, highlighting the significant changes in glycolytic metabolism involving 15 genes and cell apoptosis pathways. The heatmap of the three DEGs shown in Figure5 E demonstrated that the surgical group exhibited significantly increased expression of glycolysis-associated genes, including *MMP*, *PKM2*, and *STAT*, compared with the other two groups. This increase was accompanied by reduced expression of glycolysis-negative regulatory factors, including FTH1 and BCL2. Simultaneously, DEGs positively associated with apoptosis, including *MASP1*, *MASP2*, and *C2H2*, were upregulated, whereas DEGs negatively associated with apoptosis, including *BZW1* and *BZW2*, were downregulated.

**Figure 5.**
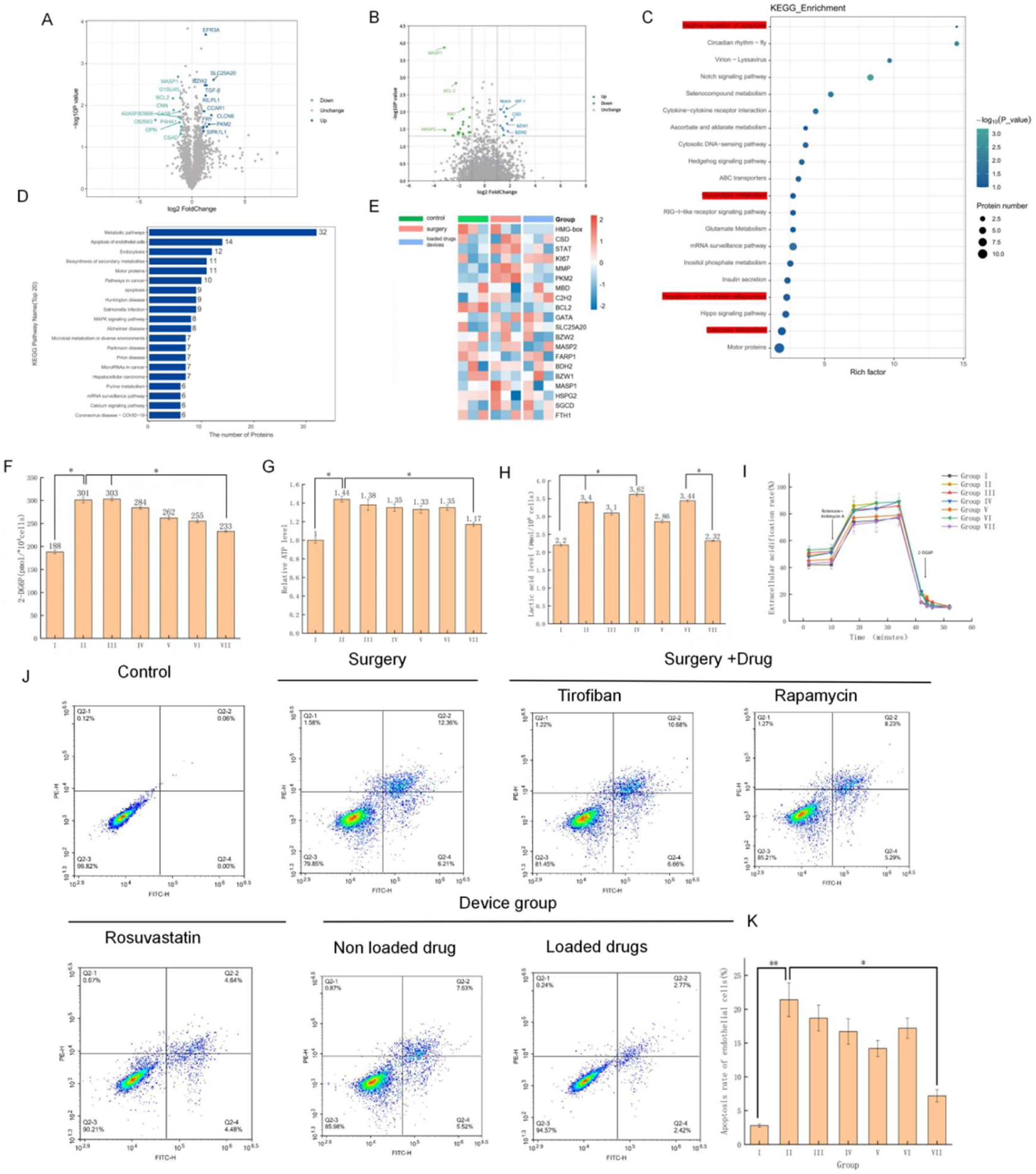
(A) RNA sequencing analysis of transplanted veins from experimental animals. (B) Volcano plot of DEGs. (C–D) KEGG enrichment analyses. (E) heatmap of DEGs. (F–I) Evaluation of glycolytic function of VSMCs in transplanted veins, including glucose uptake (F), lactate production (G), ATP production (H), and ECAR analysis (I). (J–K) Endothelial cell apoptosis assay results and quantitative analysis.

Based on these findings, glycolytic metabolism and apoptosis were further evaluated in VSMCs from each group. This analysis aimed not only to verify the roles of these pathways in restenosis but also to assess the effects of the drug-loaded microneedle device on mid-term restenosis. The results related to glycolytic metabolism in VSMCs are shown in Figures 5 F–I and include measurements of glucose uptake (Figure5 F), lactate production (Figure5 G), ATP production (Figure5 H), and extracellular acidification rate (ECAR) analysis (Figure5 I) in each group. Among multiple glycolytic metabolism indicators, Group II showed significant differences compared with Groups I and VII, confirming the association between glycolytic metabolism and mid-term restenosis in transplanted veins. These findings suggest that the drug-loaded microneedle device can effectively regulate the progression of restenosis.

The results of apoptosis experiments are shown in Figures 5 J and 6K. The apoptosis rate of cells in Group II was significantly higher than that in Groups I and IV, and the difference was statistically significant. This finding indicates that apoptosis is closely related to mid-stage restenosis in transplanted veins. Abnormal apoptosis of normal VSMCs may contribute to dysregulated cell proliferation and restenosis. The drug-loaded microneedle device may therefore help control restenosis progression by modulating these cellular processes.

**Figure 6.**
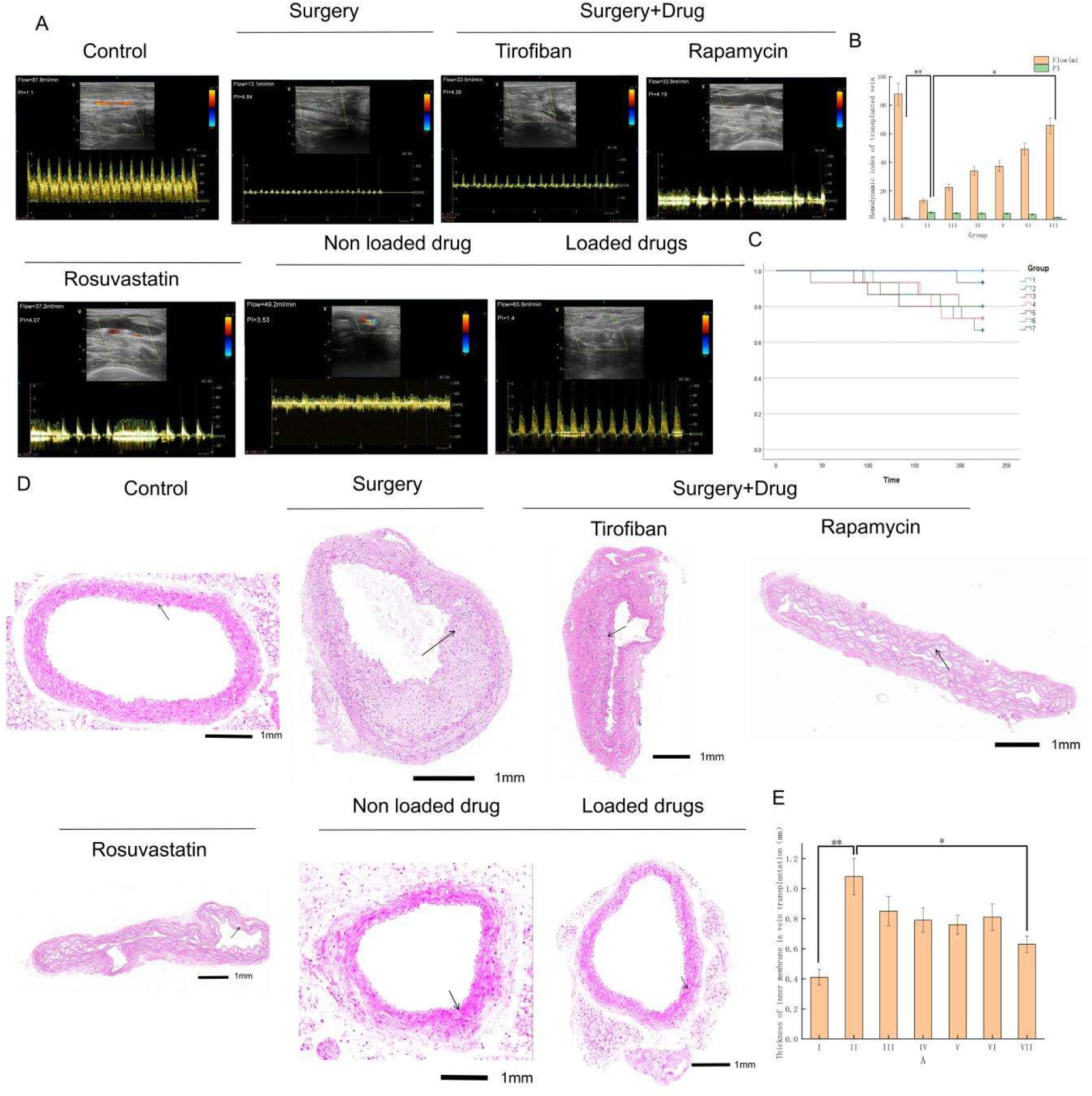
(A–B) Hemodynamic parameters and statistical analyses of transplanted veins in experimental animals at 32 weeks after surgery. (C) survival analysis curve of experimental animals. (D–E) Representative HE staining images and quantitative analysis of grafted vein intimal thickness in each group.

### 2.6 Effect of the Drug-Loaded Microneedle Device on Long-Term Restenosis of Transplanted Veins

The survival curves are shown in Figure 6C. The mortality rate in Group II was higher than that in Groups I and VII, whereas no significant differences were observed among the remaining groups. These findings indicate that long-term restenosis has a substantial impact on patient prognosis.

To evaluate the patency of transplanted veins in the long term, we performed ultrasound examinations to assess the hemodynamic parameters of transplanted blood vessels in experimental animals. Representative ultrasound images and corresponding statistical analyses are shown in Figures 6A and 6B. The blood flow rates of transplanted blood vessels in Groups I–VII were 87.8, 13.1, 22.5, 33.9, 37.2, 49.2, and 65.8 mL/min, respectively. The PI values for Groups I–VII were 1.1, 4.84, 4.39, 4.19, 4.07, 3.53, and 1.4, respectively. The hemodynamic parameters of Group II were significantly worse than those observed in Groups I and VII. Although Groups IV and V exhibited some degree of improvement in hemodynamic parameters, the effects were limited. These findings indicate that peripheral administration of therapeutic agents can improve the hemodynamic parameters of transplanted veins, but the effect is not significant. By contrast, the drug-loaded microneedle device significantly improved the long-term hemodynamic indicators of transplanted vessels, further highlighting the advantage of continuous and targeted drug release from the device. Next, we performed HE staining to evaluate the intimal thickness of transplanted veins and assess the degree of lumen compression. Representative HE staining images are shown in Figure 6D, and quantitative measurements of graft vein intima thickness are illustrated in Figure 6E. The intimal thicknesses of blood vessels in Groups I–VII were 0.41, 1.08, 0.82, 0.78, 0.81, 0.76, and 0.57 mm, respectively. Compared with Groups I and VII, Group II exhibited significantly increased vascular intima thickness, and the difference was statistically significant. These results were consistent with the observed hemodynamic results and suggest that inhibition of SMC proliferation and migration plays an important role in suppressing long-term restenosis. Although the drug-loaded microneedle device was unable to completely restore intimal thickness to levels comparable with nonsurgical conditions, the device achieved substantial therapeutic effects through sustained and targeted drug release.

### 2.7 Molecular Mechanisms of the Drug-Loaded Microneedle Device Intervention in Long-Term Restenosis of Transplanted Veins

At postoperative week 32, VSMCs were isolated from transplanted veins in each experimental group. VSMC proliferation and migration were evaluated separately using EdU and scratch assays, respectively. The EdU assay results are shown in Figures 7A and B. Compared with Groups I and VII, both the number of nuclei and proliferating cells in Group II were significantly increased, and the difference was statistically significant. The scratch assay results are shown in Figures 7C and D and demonstrated trends similar to those observed in the EdU assay. Compared with Groups I and VII, the migration level of cells in Group II was significantly increased, and the difference was statistically significant. These findings indicate that the drug-loaded microneedle device effectively controls the proliferation and migration of VSMCs derived from transplanted veins.

**Figures 7.**
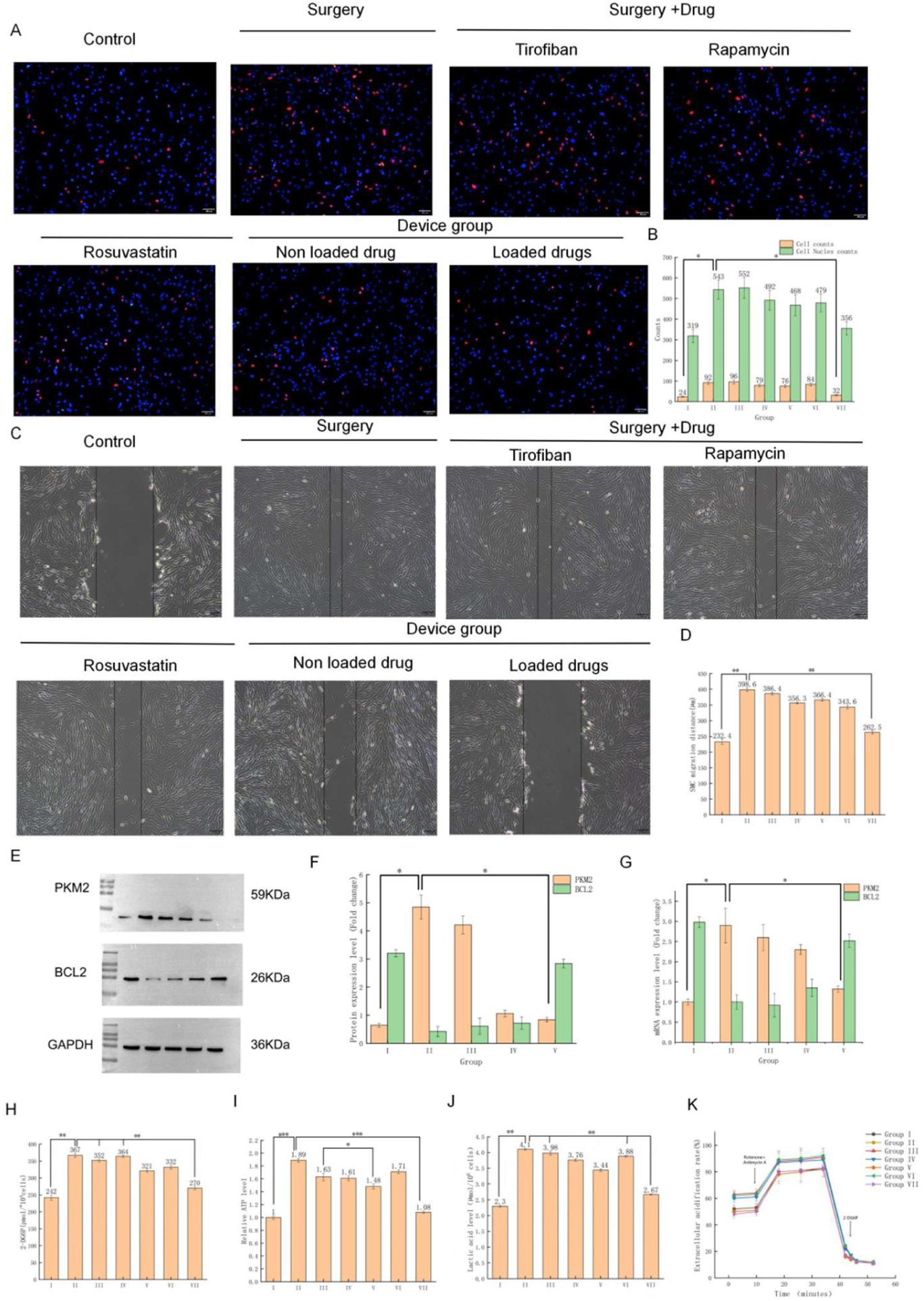
(A–B) EdU assay results and quantitative analysis of VSMCs isolated from transplanted veins at 32 weeks after surgery. (C–D) Scratch assay result and quantitative analysis of VSMCs obtained from transplanted veins 32 weeks after surgery. (E–F) Western blot results and protein expression analysis of glycolysis-related genes in transplanted veins, (G) qPCR results and mRNA expression analysis of glycolysis-related genes. (H–K) Evaluation of glycolysis activity in VSMCs obtained from transplanted veins, including glucose intake (H), ATP production (I), lactate production (J), and ECAR analysis (K).

As mentioned earlier, long-term restenosis in transplanted veins is associated with alterations in cellular glycolysis and lipid metabolism, whereas the PKM2–BCL2 pathway is closely related to cellular glycolysis. Therefore, relevant molecular indicators were further evaluated, and the Western blot and qPCR results are shown in Figures 7E–G. Protein and mRNA levels of PKM2 were significantly higher in Group II than in Groups I and VII. By contrast, protein and mRNA levels of BCL2 were significantly lower in Group II than in Groups I and VII.

To further evaluate cellular glycolysis activity, we analyzed the changes in glucose uptake (Figure 7H), ATP production (Figure 7I), lactate levels (Figure 7J), and ECAR (Figure 7K) in each experimental group. The findings suggested that the drug-loaded microneedle device may suppress the long-term restenosis of transplanted veins by inhibiting excessive glycolytic activity in VSMCs.

To further investigate the molecular mechanisms associated with long-term restenosis, we performed RNA sequencing on experimental animals in the control and surgical groups, with three animals included in each group. As shown in the volcano plot in Figure 8A, a total of 213 DEGs were identified between the control and surgical groups. The KEGG pathway enrichment analysis results shown in Figures 8B and 8C demonstrated significant changes in glycolytic metabolism involving 23 genes and endothelial cell apoptosis pathways involving seven genes. As shown in Figure 8D, heatmap analysis of the three DEGs showed that DEGs related to glycolytic metabolism, including *MMP* and *PKM2*, were significantly upregulated in the surgical group. This increase was accompanied by reduced expression of negative regulatory factors of glycolysis, including *FTH1* and *ZBTB*. Meanwhile, genes positively correlated with endothelial cell apoptosis, including *OPN* and *Ki67*, were upregulated, whereas DEGs negatively associated with apoptosis, including *CNN*, *CSB*, *α-SMA*, and *SM22-α*, were downregulated.

**Figure 8.**
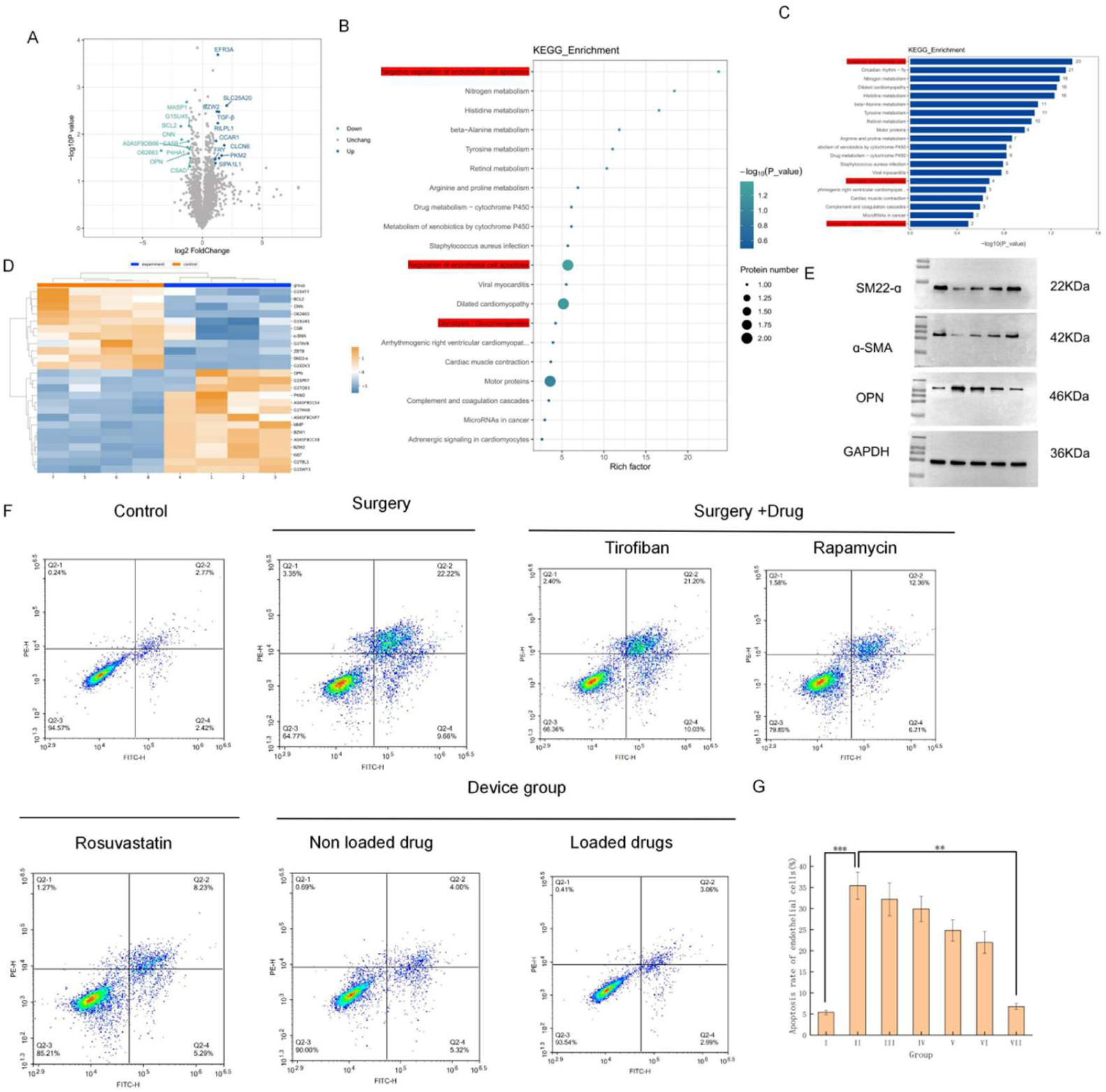
(A) Volcano plot of restenosis-related genes identified through RNA sequencing. (B–C) KEGG pathway enrichment analysis. (D) Heatmap analysis of selected genes. (E) Western bolt analysis of selected genes. (F–I) Endothelial cell apoptosis results and quantitative analyses in each group.

Western blot analysis was subsequently performed to validate the correlation of selected key genes, as shown in Figure 8E. Among these markers, *α-SMA* and *SM22-α* were identified as protective factors against endothelial cell apoptosis, whereas *OPN* was identified as a risk factor for apoptosis. The cell apoptosis results shown in Figures 8F and 8G demonstrated that endothelial cell apoptosis levels in Group II were considerably higher than those in Groups I and VII, and the difference was statistically significant.

Collectively, these findings suggest that mid- to long-term restenosis of transplanted veins is associated with endothelial cell apoptosis and excessive activation of glycolytic pathways in VSMCs, which subsequently promote excessive proliferation and migration. These mechanisms appear to represent major risk factors contributing to restenosis. The drug-loaded microneedle device targets these two pathological processes through sustained and controlled release of drugs for prolonged periods, thereby protecting transplanted veins from mid- to long-term restenosis and significantly reducing long-term postoperative mortality in experimental animals.

## 3. Discussion

In this study, we successfully developed a drug-loaded microneedle patch device capable of sequential and controlled release of three therapeutic agents—tirofiban, rapamycin, and rosuvastatin—to address the full spectrum of pathological processes involved in venous graft restenosis following CABG. The device demonstrated excellent physicochemical characteristics, including uniform pyramidal microneedles (∼700–800 μm), nanoparticle size (∼125 nm), and sufficient mechanical strength (1.07 N/needle) to penetrate the venous intima without cytotoxicity or hemolysis (Figure 1). Using a mouse model of venous bypass grafting involving inferior vena cava-to-carotid artery interposition, we demonstrated that the developed microneedle patch significantly improved graft patency at acute (72 h), mid-term (4 weeks), and long-term (32 weeks) stages, as assessed through ultrasonography (blood flow velocity and PI), histomorphometric analyses (intimal thickness), and molecular investigations (Western blotting, qPCR, and RNA sequencing). Notably, our findings suggest that excessive glycolysis-driven SMC proliferation, coupled with dysregulated SMC apoptosis, represents a key mechanism contributing to mid- to long-term restenosis. To the best of our knowledge, this is the first study to integrate perivascular mechanical support with sequential triple-drug release to target vein graft failure across all pathological stages.

### 3.1 Distinct Mechanisms at Different Restenosis Stages

#### 3.1.1 Early Phase (Perioperative Thrombosis; 0–7 Days)

The LC-MS/MS release profile showed that tirofiban, a GP IIb/IIIa antagonist, was released rapidly (∼37% within 3 days and 65% within 7 days), matching the critical time window for platelet thrombus formation. In the surgical control group (Group II), platelet thrombus incidence reached 23.3% at 72 h and was accompanied by elevated platelet counts and prolonged APTT/PT parameters. By contrast, the microneedle patch–treated group (Group VII) showed substantially reduced thrombus incidence (10%), improved coagulation parameters, and significantly reduced GP IIb/IIIa protein expression (Figures 2D–3F). Although systemic tirofiban administration (Group III) also inhibited thrombosis, previous clinical studies^[17–19]^ reported an increased risk of major bleeding in humans. Localized perivascular delivery through the microneedle patch minimized systemic exposure while achieving effective drug distribution within the vascular wall, as confirmed by immunofluorescence analysis (Figure 1H).

#### 3.1.2 Mid-Term Phase (Intimal Hyperplasia; 1–8 Weeks)

Rapamycin release commenced approximately on day 12 (12% by day 14 and 54% by week 4), corresponding to the period during which SMC activation and phenotypic switching became prominent. At 4 weeks, Group VII exhibited significantly improved blood flow (98.1 vs. 46.4 mL/min in Group II) and reduced PI (1.4 vs. 5.2 in Group II), accompanied by markedly decreased intimal thickness (0.45 vs. 0.84 mm in Group II) (Figure 3). EdU and scratch assays further confirmed that the microneedle patch suppressed SMC proliferation and migration more effectively than systemic rapamycin administration (Figures 4A–5D).

Mechanistically, we identified HIF-1α as a key transcriptional regulator: Group II showed significantly elevated HIF-1α protein and mRNA expression levels, whereas these alterations were normalized in Group VII (Figures 4E–5G). Furthermore, PKM2 (a glycolytic enzyme) expression was upregulated and BCL2 (an anti-apoptotic marker) expression was downregulated in Group II, indicating a metabolic shift toward glycolysis and increased susceptibility to apoptosis (Figures 4H–5J).

#### 3.1.3 Late Phase (Atherosclerosis; >8 Weeks)

Rosuvastatin release began at day 40 (58% at 20 weeks and 91% at 35 weeks), matching the expected onset of graft atherosclerosis. At 32 weeks, Group VII demonstrated superior outcomes compared with the other intervention groups, indicating increased blood flow (65.8 vs. 13.1 mL/min in Group II) and reduced intimal thickness (0.57 vs. 1.08 mm in Group II) (Figure 6). Furthermore, survival curve analysis revealed increased mortality in Group II, underscoring the clinical relevance of maintaining long-term graft patency.

### 3.2 A novel Mechanistic Insight: Glycolysis–Apoptosis Coupling in SMCs

RNA sequencing analysis (Figure5) identified 154 DEGs between the control and surgical groups, with significant enrichment in glycolytic metabolism (15 genes) and apoptosis-related pathways. Heatmap analysis demonstrated the upregulation of pro-glycolytic genes (including *MMP*, *PKM2*, and *STAT*) and downregulation of anti-glycolytic/anti-apoptotic genes (such as *FTH1* and *BCL2*) in Group II compared with Group VII. Importantly, we directly measured functional glycolytic activity in isolated SMCs. Group II exhibited significantly increased glucose uptake, lactate production, ATP generation, and ECAR, all of which were restored to baseline levels in Group VII (Figures 5 F–6I). Simultaneously, flow cytometry revealed that the SMC apoptosis rates were significantly increased in Group II, which paradoxically contributed to restenosis by creating a pro-inflammatory, pro-proliferative microenvironment (Figures 5 J–6K).

Based on these findings, we propose a potential pathological cycle in which mechanical injury and abrupt pressure alterations activate HIF-1α signaling, resulting in PKM2-mediated enhancement of aerobic glycolysis in SMCs (the Warburg effect). Enhanced glycolytic activity promotes excessive SMC proliferation and migration, while simultaneous mitochondrial dysfunction and reduced BCL2 expression may contribute to apoptosis in SMCs. Apoptotic pathways further promote survival signaling in neighboring SMCs, thereby perpetuating intimal hyperplasia. This “glycolysis-apoptosis coupling” model provides a novel framework for understanding the development of mid-term restenosis and suggests that dual targeting of metabolic reprogramming and cell survival pathways may be more effective than monotherapies.

### 3.3 Clinical Relevance and Translational Potential

From a clinical perspective, the proposed device addresses several unmet needs. First, current guidelines recommend initiation of dual antiplatelet therapy within 4–24 h after CABG; however, prolonged postoperative mechanical ventilation frequently delays treatment initiation. Localized tirofiban release from the microneedle patch may help bridge this therapeutic gap without increasing bleeding risk. Second, systemic rapamycin and statin administration have shown limited efficacy in preventing vein graft failure owing to poor local penetration. The microneedle patch achieved sustained, high local concentrations. Third, the mechanical support provided by the hydrogel-based microneedle patch itself may provide mechanical support by limiting acute vein overexpansion, thereby complementing its pharmacological effects.

To our knowledge, no currently available material or device simultaneously targets thrombotic, hyperplastic, and atherosclerotic processes involved in vein graft disease. Therefore, our work completes the research, development, and pilot production of a device with independent intellectual property, offering a novel strategy for improving outcomes following CABG.

### 3.4 Limitations and Future Directions

Several limitations should be acknowledged. First, although the mouse cuff model is widely used for mechanistic investigations, murine vessels differ substantially from human saphenous veins in terms of vessel size, wall structure, and hemodynamic forces. Therefore, future studies should validate device efficacy in larger animal models, such as rabbit or porcine jugular vein-to-carotid artery interposition models, prior to clinical translation. Second, while RNA sequencing revealed multiple molecular alterations, we did not perform single-cell sequencing to distinguish the contributions of SMCs, endothelial cells, and immune cells. Third, although the 32-week data were encouraging, longer follow-up periods extending to 1–2 years are required to fully evaluate atherosclerosis progression in a manner comparable to human disease. Finally, biomechanical factors such as cyclic stretching and shear stress, which may initiate early SMC activation, warrant further investigation using in vitro mechanical stimulation systems. Future research should also include clinical feasibility studies with histological analyses of retrieved human graft samples to validate drug distribution and target engagement.

Current ongoing investigations include: (1) a prospective clinical registry evaluating vein graft patency in patients receiving similar perivascular devices; (2) mechanistic validation studies using CRISPR-based manipulation of HIF-1α and PKM2 in SMCs; and (3) a large-animal study evaluating long-term safety and efficacy.

Despite these limitations, the present study provides proof-of-concept evidence supporting a first-in-class integrated device that targets vein graft restenosis across pathological stages and may provide a novel strategy for perioperative vascular protection.

## 4. Conclusion

In conclusion, to address the full spectrum of pathological mechanisms involved in venous graft failure following CABG, including early thrombosis, mid-term intimal hyperplasia, and late atherosclerosis, this study successfully designed and validated a perivascular drug-loaded microneedle patch device capable of sequential and controlled release of tirofiban, rapamycin, and rosuvastatin. In a mouse model of venous bypass grafting, the device significantly reduced perioperative platelet thrombus formation through local GPIIb/IIIa inhibition, alleviated mid-term intimal hyperplasia through sustained delivery of rapamycin targeting HIF-1α/PKM2-mediated glycolysis, and improved long-term graft through rosuvastatin release, ultimately reducing long-term mortality. Mechanistically, we identified a novel “SMC cell glycolysis–proliferation/migration–endothelial cell apoptosis coupling” axis as a key driver of mid- to long-term restenosis.

Our work addresses a key clinical bottleneck in the surgical management of coronary heart disease by providing a safe, effective, and locally targeted intervention that reduces systemic bleeding risks associated with conventional antiplatelet therapy while simultaneously targeting the mechanisms underlying mid- to long-term restenosis. The successful development of this microneedle patch may provide a novel approach for perioperative vascular protection and has the potential to improve long-term prognosis in patients undergoing CABG.

Future research should focus on single-cell sequencing to characterize cellular heterogeneity within the graft microenvironment and on biomechanical investigations to elucidate stretch-induced SMC activation mechanisms. In addition, large-animal studies and subsequent clinical trials are required to further establish the safety and efficacy of this device in human applications.

## 5. Experimental Section/Methods

### 5.1 Main materials and equipment

#### 5.1.1 Main materials

Hyaluronic acid (HA) was obtained from Shandong Huaxi Biotechnology Co., Ltd. Tirofiban (purity: 98.5%) was purchased from Sigma Aldrich (Shanghai) Trading Co., Ltd., China. Rapamycin (purity: 98%) was purchased from Bailingwei Technology Co., Ltd. Rosuvastatin (ZD4522, purity: 98%) was purchased from Medchemexpress Co., Ltd. (MCE, China). PDMS templates were obtained from Henan Weina Pentium Biotechnology Co., Ltd. Deionized water was prepared using a Millipore purification system. Acetone (chromatographic grade) was purchased from Bailingwei Technology Co., Ltd.

All animal experiments were conducted in strict accordance with national regulations and guidelines for animal welfare and ethics and were approved by the Animal Ethics and Welfare Committee of Beijing Anzhen Hospital Affiliated to Capital Medical University.

A total of 210 C57BL/6 wild-type (WT) mice were obtained from Beijing Anzhen Animal Center affiliated with Capital Medical University and randomly assigned to seven groups: Group I, control group; Group II, surgical arteriovenous transplantation group; Groups III–V, surgery groups receiving continuous peripheral administration of tirofiban, rapamycin, and rosuvastatin, respectively; Group VI, non-drug-loaded microneedle device group; and Group VII, drug-loaded microneedle device group.

#### 5.1.2 Main Instruments and Equipment

The following instruments and equipment were used in this study: an electric blast drying oven (Shanghai Jiecheng Experimental Instrument Co., Ltd.); a Fourier transform infrared spectrometer (Invenio, Bruker, Germany); field-emission scanning electron microscope (JSM-IT800SHL, Nippon Electronics); LC–MS/MS (AB SCIEX qtrap™ 6500), a TEM (Hitachi ht7800); a laser particle sizer (NanoZS, Malvern); a GT100X vibration ball mill (Beijing gredeman Instrument Equipment Co., Ltd.); a digital scanner (Pannoramic MIDI, 3DHISTECH, Hungary); and TMS circular stretching culture system (ATMS Boxer, Taiwan).

### 5.2 Methods

#### 5.2.1 Preparation of the Drug-Loaded Microneedle Patch Device

A drug-loaded nanoparticle hydrogel solution was prepared for the controlled release of the three therapeutic agents. The preparation procedure for the controlled-release drug-loaded nanoparticle hydrogel solution has been previously reported in *Advanced Science*^[24]^ and is therefore not described in detail here.

First, 0.2 g of HA was added to 20 mL of deionized water and mixed at room temperature using a mechanical stirrer until a gel-like homogeneous solution was obtained. The prepared solution was added to the HA solution and mixed thoroughly to ensure uniform dispersion. The resulting drug-loaded hydrogel mixture was then transferred onto a PDMS mold and placed in a vacuum drying oven. Vacuum treatment was performed at room temperature under a pressure of -0.1 MPa for approximately 10 min.

Because vacuum treatment may cause partial overflow of the hydrogel from the mold, additional HA hydrogel solution was added to refill the PMDS mold as needed, followed by repeated vacuum treatment until no visible air bubbles remained within the mold structure. After complete drying, the mold was removed, and the drug-loaded microneedle protection device was obtained. The preparation process is illustrated in Figure S1A.

#### 5.2.2 Physical Properties of the Drug-Loaded Microneedle Device

##### Morphological Characterization of Microneedle Patch Morphology

The prepared drug-loaded microneedle device was cut into square sections measuring 1.5 cm × 1.5 cm. The microneedle patch was fixed using a sample holder, and microneedle morphology was observed using SEM. Microneedle height, bottom diameter, and structural integrity were measured to evaluate the physical and mechanical properties of the device.

##### Evaluation of Drug Incorporation Using Infrared Spectroscopy

Drug-loaded microneedle device samples were collected and placed in a mortar. Dry potassium chloride powder was added as a dispersing agent at a mass ratio of approximately 100:1 relative to the sample and was thoroughly ground and mixed. The resulting mixture was transferred into a compression mold and compressed into transparent test specimens. Infrared spectra of the samples were then collected using a Fourier transform infrared spectrometer to evaluate whether the three drugs had been successfully incorporated into the microneedle patches.

##### Drug Release Analysis of Drug-Loaded Microneedle Patches Using LC–MS/MS

To preliminarily evaluate the drug release ability of drug-loaded microneedles in target organs, we established a method based on high-performance LC–MS for the quantitative measurement of the three drugs in biological samples. This method was used to continuously monitor drug release from the microneedles into simulated biological environments at different times.

For chromatographic analysis, a Waters XBridge BEH C18 column (2.1 mm × 50 mm, 3.0 μm) was used. The mobile phase consisted of pure water containing 0.1% formic acid (phase A) and pure methanol (phase B). The flow rate was set to 0.4 mL/min, the column temperature was maintained at 40 ℃, and the injection volume was 5 μL. Gradient elution conditions are listed in Table 3. Mass spectrometry conditions were as follows: electrospray ionization in positive ion mode and multiple reaction monitoring scanning. Tirofiban, rapamycin, rosuvastatin, and carbamazepine (internal standard) were analyzed. The detected ion pairs were m/z 652.4 → m/z 277.6, m/z 852.6 → m/z 599.3, m/z 922.8 → m/z 426.3, and m/z 237.0 → m/z 192.0, respectively. The declustering potential was 80eV, the collision voltage was 70eV, the atomization temperature was 450 ℃, and the source temperature was 100 ℃.

| TIME (min) | FLOW (mL/min) | Mobile<br>phaseA (%) | Mobile<br>phaseB (%) |
| --- | --- | --- | --- |
| 0 | 0.4 | 80.0 | 20 |
| 0.2 | 0.4 | 80.0 | 20 |
| 0.8 | 0.4 | 10.0 | 90 |
| 3.2 | 0.4 | 10.0 | 90 |
| 3.21 | 0.4 | 80.0 | 20 |
| 6.0 | 0.4 | 80.0 | 20 |

##### Vascular-Targeted Drug Release Study Using Discarded Human Great Saphenous Vein Samples

A vascular-targeted drug release study was conducted using discarded great saphenous vein tissue obtained from patients undergoing CABG surgery. The experimental protocol was approved by the Medical Ethics Committee of Beijing Anzhen Hospital (approval number: KS2022051)This study obtained informed consent from patients and was conducted in accordance with the principles of the Helsinki Declaration. The vessel samples were rinsed with physiological saline and cut into sections measuring 1.5 cm × 1.5 cm. Microneedle patches (1.5 cm × 1.5 cm) were applied to vascular tissue and wrapped in moist sterile gauze to maintain hydration. Tissue samples were collected at 1, 3, 6, 12, and 24 h. At each time point, vascular tissues were washed with physiological saline, residual surface liquid was removed, and the tissues were weighed and cut into small pieces. Physiological saline (1 mL) was added for tissue homogenization at a tissue-to-saline ratio of 1:5. Homogenization was performed at 60 Hz for 60 s. After homogenization, samples were stored in a -20 ℃ until further analysis.

##### Determination of Drug Release in Tissue Samples

Drug concentrations within vascular tissues were measured using a protein precipitation method. Briefly, 50 µL of vascular tissue homogenate was transferred into an anti-adsorption EP tube. Subsequently, 50 µL of internal standard working solution containing carbamazepine (25 µL/mL) and 0.9 mL of methanol containing 2% formic acid were sequentially added to precipitate the protein. The mixture was vortexed for 1 min and centrifuged at 4 °C for 5 min at 13,300 rpm. Subsequently, 20 µL of the supernatant was collected for LC–MS/MS analysis, and the concentration of each drug was measured at different time points. This study evaluated the drug release profiles for up to 1 year to evaluate the long-term controlled release characteristics of the device throughout the entire restenosis process.

##### Mechanical Property Evaluation Using an Electronic Tensile Testing Machine

According to our previous research,^[20]^ drug-loaded microneedles intended to penetrate the venous intima must withstand a force of 0.5 N applied by a pressure plate. Therefore, the speed at which the pressure plate moved toward the microneedle was set to 0.5 mm/s to determine whether the microneedle could tolerate and generate sufficient force to penetrate the vascular intima.

##### Evaluation of Drug Distribution in the Venous Intima Using Immunofluorescence

Residual unused human great saphenous vein tissue obtained during CABG surgery (provided by Anzhen Hospital) was immersed in 0.9% physiological saline at room temperature and gently dried with sterile gauze before use. We prepared a drug-loaded microneedle device using rhodamine B-labeled tirofiban, CY5-labeled rapamycin, and CY7-labeled rosuvastatin. The microneedle patches were evenly wrapped around the outer membrane of the great saphenous vein and gently pressed to ensure insertion; after this procedure, the patch was removed.

After 10 min, the microneedles were allowed to completely dissolve, and a 5-mL syringe was used to inject water into the distal end of the vein to determine whether vein leakage occurred and to preliminarily assess whether microneedle insertion caused vascular damage.

The treated great saphenous veins were immediately placed in cryogenic tubes, stored in liquid nitrogen, and frozen for 48 h. Samples were embedded using optimal cutting temperature compound, and frozen sections were prepared. Immunofluorescence imaging was subsequently performed using a digital scanning system at different time points to evaluate drug distribution within the outer and inner layers of the great saphenous vein. Drug distribution patterns were further used to assess the degree of microneedle patch penetration into the great saphenous vein.

#### 4.2.3 Biosafety Assessment of the Drug-Loaded Microneedle Device

Because hemolysis may lead to severe adverse biological effects, hemolysis testing was performed to determine the biocompatibility of the drug-loaded microneedle device and the developed formulations. Materials exhibiting hemolysis rates below 5% were considered non-toxic.^[35]^ The experiment consisted of four groups: a negative control group, a hydrogel solution treatment group, a drug-loaded hydrogel solution treatment group, and a positive control group (Groups I–IV, respectively). Blood samples were collected from healthy human volunteers and centrifuged at 825 × g for 15 min to isolate red blood cells (RBCs). The supernatant was discarded, and RBCs were washed three times with phosphate-buffered saline (PBS). Subsequently, a 2% (v/v) RBC–PBS solution was prepared. Briefly, 100 µL of the RBC suspension was mixed with 100 µL of the drug-loaded formulation solution (1 mg/mL RUT concentration) and incubated at 37 °C for 1 h using physiological saline as the diluent. Following incubation, the mixture was centrifuged at 825 × g for 5 min. Subsequently, 200 μL of the supernatant was transferred into a 96-well plate. PBS (0.01M, pH 7.4) and 1% Triton X were used as negative and positive controls, respectively, and equal volumes of RBC suspension (100 µL) were added to each control group. Absorbance was measured at 540 nm using a microplate reader. The hemolysis rate was calculated using the following equation^[36]^:

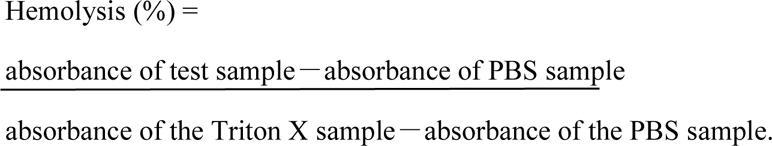

##### In Vitro Cytotoxicity Assessment of the Drug-Loaded Microneedle Device

HUVSMCs were used to evaluate the cytotoxicity of the drug-loaded microneedle device, and dimethyl sulfoxide (DMSO) was used as a solvent control. Drug-loaded microneedle device solutions at different concentrations were used as experimental groups. HUVSMCs were cultured in growth medium and maintained in an incubator at 37 °C with 5% CO_2_. Cells were observed under a microscope and passaged when cell confluence reached approximately 80– 90%. This study was approved by the Ethics Review Committee of Beijing Anzhen Hospital (approval number: KS2022051).

For the Cell Counting Kit-8 (CCK-8) assay, a blank control group, DSMO solvent control group, and drug-loaded microneedle device solution group with different concentrations were prepared. Five milliliters of each drug-loaded hydrogel solution was transferred into sterile centrifuge tubes, followed by the addition of 1 mL of DMSO. Samples were dissolved using ultrasonic treatment.

HUVSMCs in the logarithmic growth phase were digested with trypsin, collected, centrifuged, and quantified under a microscope to obtain a cell suspension at a concentration of 3 × 10^4^ cells/mL. Subsequently, 100 µL of cell suspension was seeded into each well of a 96-well plate, with four replicate wells included per group. Wells containing 100 µL of blank culture medium served as controls. Cells were incubated at 37 °C for 24 h to allow attachment. During the inoculation process, peripheral wells were avoided and filled with PBS to prevent evaporation of the culture medium and reduce edge effects.

Following cell adhesion, each group was treated with the corresponding drug solution. Drug concentrations used in the experimental group are described in the corresponding section. Equivalent volumes of DMSO were added to the solvent control group. Cells were treated for 12, 24, 48, and 72 h, with measurements obtained at baseline (0 h) and at 12, 24, and 72 h after treatment.

Before CCK-8 measurements, the culture medium was replaced to eliminate the interference of drug formulations with absorption measurements. Subsequently, 10 µm of CCK-8 reagent was added to each well, and the plate was incubated at 37 °C in a 5% CO_2_ incubator for 3 h. Absorbance was measured at 450 nm using a microplate reader, and the absorbance values for each well were recorded.

#### 5.2.4 Establishment of Animal Models and Evaluation of Postoperative Grafted Vessel Quality

A mouse vein graft model was established using the “cuff” technique, which mimics key features of human CABG surgery. Briefly, the inferior vena cava was harvested from donor mice, and the venous end was sutured to an arterial cuff, followed by transplantation of the inferior vena cava into the right internal carotid artery of recipient mice. During surgery, a ×15 operating microscope was used for magnification, and vascular anastomosis was performed using 10-0 Prolene sutures. Complete vascular anastomosis was confirmed before closure.

The diameter and stenosis rate of the transplanted veins were evaluated using ultrasonography. Blood flow measurements were obtained at 24 h, 72 h, 4 weeks, 16 weeks, and 32 weeks after surgery.

The vascular lumen loss rate was calculated using the following formula:

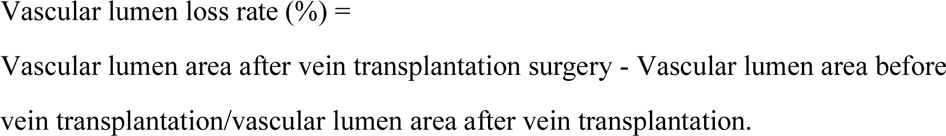

Blood flow and PI were used to evaluate vascular function and resistance to blood flow within the transplanted blood vessels and were assessed using ultrasonography. The PI was calculated as follows:

PI = PSV-MDV/MV,

where PSV is the peak systolic velocity, MDV denotes the mean diastolic velocity, and MV is the mean velocity.

At 24 h, 72 h, 4 weeks, 16 weeks, and 32 weeks after surgery, the mice were euthanized.Euthanasia of mice by injection of pentobarbital sodium(200mg/kg), All procedures were performed by trained personnel following standard protocols, with personal protective equipment worn throughout,Carcasses were placed in biohazard bags and incinerated according to institutional guidelinesand HE staining was performed to evaluate the degree of intimal hyperplasia in transplanted blood vessels from each group. Before tissue collection, physiological saline containing heparin was perfused through the circulation to remove residual blood. Jugular vein graft samples were subsequently harvested, fixed in 10% formalin, dehydrated, and embedded in paraffin. Tissue samples were sectioned into 4-µm-thick slices and stained using HE. Histological images were collected and analyzed using a NIKON NIS-Elements imaging system.

Intimal hyperplasia was quantified by measuring intimal thickness. On each tissue slice, 16 points surrounding each vein graft cross-section were randomly selected, and intimal thickness was measured at each point. The average thickness for each graft was then calculated. Three tissue sections were obtained from each rabbit, and the mean value from the three tissue cross-sections was used as the representative intimal thickness measurement.

#### 5.2.5 Molecular Mechanism Study of the Drug-Loaded Microneedle Device on Early Restenosis (Perioperative Thrombosis) of Transplanted Veins

Peripheral blood samples were collected from experimental animals on postoperative days 1, 2, 3, 7, 10, and 14 following intravenous transplantation. Samples were collected into anticoagulant tubes, mixed thoroughly, and stored at 4 °C for no longer than 30 min prior to processing. Whole blood samples were centrifuged at 3,000 × g for 15 min at 4 °C, and the supernatants were transferred into labeled sample tubes for storage until subsequent analysis.

After sample preparation, serum samples were divided into two portions. The first portion was used to determine the platelet count, PT, and APTT. The second portion was used to evaluate the effects of the treatment group (II, IV, and V), matrix hydrogel solution. and microneedle drug delivery system on GP IIb/IIIa receptor expression in the peripheral blood using enzyme-linked immunosorbent assay (ELISA) kits (Shanghai Yanji Biotechnology Co., Ltd., China). Briefly, GP IIb/IIIa proteins in the standards and samples bound to monoclonal antibodies. Biotinylated anti-human GP IIb/IIIa antibodies were added to form immune complexes attached to the plate surface. Horseradish peroxidase-labeled streptomycin was then added to bind biotin, followed by substrate solution, resulting in a colorimetric reaction (blue). Finally, sulfuric acid was added to terminate the reaction, and OD values were measured at 450 nm. The concentrations of GP IIb/IIIa were directly proportional to OD values and were calculated according to standard curves. Western blotting was subsequently performed to verify GP IIb/IIIa protein expression levels in each group.

For molecular biology experiments, we extracted VSMCs and endothelial cells from transplanted veins harvested from experimental animals. Briefly, animals were euthanized at designated time points, and transplanted vein samples were collected. VSMCs were isolated using a combination of enzymatic digestion and tissue block adhesion methods. Clean graft veinswere placed into enzyme digestion solution and incubated in a constant-temperature shaker at 37 °C for 45 min, depending on vessel size and enzymatic activity. Digestion was terminated when the blood vessels became translucent, and the surrounding tissues exhibited flocculent structures. The outer, inner, and intermediate membrane layers of the blood vessels were carefully separated.

The isolated membrane layers were washed with Hank’s Balanced Salt Solution containing Ca^2+^ and Mg^2+^ and cut into 1–2-mm^3^ pieces in sterile culture dishes. Tissue fragments were transferred to six-well plates containing a complex enzyme solution for secondary digestion and incubated for approximately 1 h with gentle shaking every 20 min. After 50 min of digestion, large numbers of cells were observed emerging from tissue fragments in the form of grapes and beginning to dissociate. At this stage, the SMC culture medium containing 10% serum was added to terminate digestion. The digestion solution was gently aspirated using a 1-mL syringe until cells and tissue fragments were separated and subsequently filtered through a 40-μm cell sieve to obtain a cell suspension. The tissue was removed from the filter, and the pumping and filtration steps were repeated; the cell suspension was passed through multiple filters and finally added into 15-mL centrifuge tubes. This suspension was centrifuged at 1,000 rpm for 5 min, and an SMC suspension containing 10% fetal bovine serum was collected and cultured in a six-well plate. The newly isolated P0 generation VSMCs and endothelial cells were suspended in an SMC culture medium (Gibco DMEM medium, thermophilic fish, China) and cultured for 2 days in a static incubator at 37 °C with 5% CO_2_. After 48 h of observation, the cells adhered to the wall and started to spread. The medium was replaced with fresh culture medium. After approximately seven days of cultivation, the cells were passaged, and the serum level dropped to 2% of the normal serum after passage.

Our previous research^[17–20]^ showed that, although early graft restenosis was not directly associated with intimal thickening, activation of VSMC proliferation and migration had already occurred. Therefore, we conducted EdU and scratch assays using extracted VSMCs. For the EdU assay, EdU reagent A (Ruibo, China) was added to the VSMCs isolated from each experimental group and incubated overnight.

Following incubation, the cells were washed two to times with PBS to remove excess EdU. Cells were fixed using 4% paraformaldehyde at room temperature for 30 min. The fixative was discarded, and 500 µL of glycine solution (2 mg/mL) was added to each well and incubated for 5 min on a decolorization shaker to neutralize excess polyformaldehyde. Glycine solution was discarded, and cells were washed three times with PBS for 5 minutes each. After washing, 500 µL of 0.5% Triton-100 solution was added to each well and incubated on a decolorization shaker for 10 min, followed by three additional PBS washes for 5 min each. Apollo staining solution was prepared according to the manufacturer’s instructions. The amount of staining solution used was related to the cell volume, and 50 µL of staining solution is sufficient for every 24 wells of glass slides. The staining solution was dropped onto labeled glass slides, and washed cell slides were inverted into the staining solution; the cell surface was allowed to come into contact with the staining solution. Avoiding light, the slides were incubated at room temperature for 30 min. After staining, the slides were washed three times with PBS for 5 min each. Following the steps described above, the cell nuclei were stained with DAPI staining solution and incubated at room temperature for 5 min. After staining, excess staining solution was washed one to two times with PBS on a decolorization shaker for 5 min. Subsequently, 10 µL of glycerol was added to clean glass slides, and the slide was inverted onto glycerol to avoid bubbles. After surgery, photographs were taken under a microscope.

The scratch assay involved using a marker pen and a ruler to evenly draw horizontal lines on the back of a six-hole plate, with one line drawn every 0.5–1 cm and five lines drawn per hole. VSMCs extracted from each group were cultured at 37 ℃ in a 5% CO_2_ incubator until the cell density reached 80–90%. After 48 h, a sterile pipette tip guided by a ruler was used to draw a straight line perpendicular to the longitudinal axis of the six-well plate. The cells were washed three times with PBS to remove scratched cells and cultured in a 1% oxygen-deficient incubator; the healing process was recorded for 48 h using an inverted microscope. The same procedures were used in subsequent experiments to evaluate the proliferation and migration of VSMCs at different experimental stages; therefore, these methods are not repeated in later sections.

#### 5.2.6 Molecular Mechanism Study of the Drug-Loaded Microneedle Device in Mid-Stage Restenosis Intervention of Transplanted Veins

The EdU and scratch assays used to evaluate VSMC proliferation and migration have been described previously and are therefore not repeated here. In our previous research, we identified HIF-1α as a potential therapeutic target for mid-term restenosis, with rapamycin as a key intervention agent. Therefore, we performed Western blot analysis to evaluate HIF-1α protein expression levels in Groups I, II, IV, VI, and VII. qPCR was additionally used to evaluate the therapeutic effects of the drug-loaded microneedle device.

HIF-1 α is also recognized as an important mediator of cellular glycolytic pathways. Previous studies on restenosis have largely suggested that glycolytic metabolism in SMCs is associated with long-term restenosis^[17–20]^, whereas limited information is available regarding glycolytic activation during mid-term restenosis. Therefore, Western blotting was performed to determine the expression levels of downstream factors PKM2 and BCL2 in Groups I, II, IV, VI, and VII. In addition, mRNA expression levels of PKM2 and BCR2 were evaluated by qPCR to preliminarily explore the potential involvement of glycolytic activation in SMCs during mid-term restenosis.

The qPCR procedure was performed as follows. Total RNA was isolated from vascular tissues using a Trizol detection kit (Life Technology, USA). RNA was reverse-transcribed into complementary DNA using an RNA reverse transcription kit (GeneCopoeia, USA). The reaction mixture was configured according to the composition described in Table S1.

Real-time fluorescence qPCR procedure was performed using a two-step protocol under the following conditions: initial denaturation at 95 ℃ for 2 min, followed by 40 amplification cycles consisting of denaturation at 95 ℃ for 15 s and annealing at 60 ℃ for 30 s. The absorbance was measured during each measurement step. GAPDH was used as the internal housekeeping gene. Detailed PCR conditions are presented in Table S2. Following amplification, melting curve analysis was performed from 65 to 95 ℃, with temperature increments of 0.5 ℃ per second. Relative mRNA expression levels of HIF-1α and housekeeping genes were analyzed. TableS3 lists the primer sequences.

To investigate intracellular signaling pathways associated with mid-term restenosis after vein transplantation, unbiased RNA sequencing analysis was performed on the aorta samples obtained from Groups II, VI, and VII.

##### Gene Ontology (GO) Functional Annotation

GO functional annotation of target proteins was performed using Blast2GO (V1.4.4).^[46]^ The GO annotation workflow consisted of four major steps: sequence alignment (Blast), GO terms extraction (Mapping), GO annotation (annotation), and supplementary annotation (ANNEX). Initially, all protein sequences were aligned to the corresponding database (determined by project) using NCBI BLAST+ (ncbi-blast-2.3.0+) on a Linux server. Only the top 10 aligned sequences with E-value ≤1 × 10^-3^ were retained. Second, GO terms associated with the aligned sequence with the highest bit score were selected using the Blast2GO Command Line (database version: go_20190701. obo). During annotation, factors including sequence similarity, reliability of GO source information, and GO-directed acyclic graph structures were considered. The selected GO terms were annotated to the target proteins using the Blast2GO Command Line.

InterProScan (interproscan-5.30-69.0)^[47]^ was used to identify conserved protein motifs within the EBI database, and motif-related functional information was incorporated into the annotation process. Finally, ANNEX was applied to improve the accuracy of the annotations and establish relationships among different GO terms.

##### KEGG Pathway Annotation

In the KEGG database, KEGG Orthology (KO) is used as a classification system for genes and gene products. Orthologous genes and their products with similar functions within the same pathway are assigned identical KO (or K) labels.

For KEGG pathway annotation, KOALA (KEGG Orthology and Links Annotation, V3.0)^[48]^ was used to align target protein sequences against the KEGG GENES database (version: KO_INFO_END.txt (2023.10.17). These protein sequences were then classified according to KO identifiers, and pathway information was automatically generated.

##### Enrichment Analysis of GO and KEGG Pathways

GO and KEGG pathway enrichment analyses were performed using consistent analytical algorithms. Based on Fisher’s exact test, we compared the distributions of individual GO terms and KEGG pathways in target and total protein datasets to evaluate the significance level of the enrichment of GO terms and KEGG pathways. Next, heatmaps were generated to identify significantly altered pathways, and qPCR analysis was performed to further validate pathway-associated genes with significant expression differences.

Subsequently, we evaluated the glycolytic activity in SMCs from each experimental group. Cellular glucose uptake was measured using a glucose oxidase method.

Cellular ATP production was measured using an ATP detection kit, and lactate levels were measured using ELISA. The ECAR was measured using an XF24 extracellular flux analyzer (Seahorse Biosciences) equipped with an Agilent Seahorse XF Glycolysis Rate Assay Kit and an XF96 extracellular flux analyzer (Seahorse Biosciences) equipped with an Agilent Hippocampus XF Mito Stress Test and Glycolysis Stress Rest Kits.

Because mid-term restenosis has been associated with endothelial cell apoptosis, apoptosis was assessed using flow cytometry. Cells were washed with Dulbecco’s phosphate-buffered saline (21 031-CV; CORNING) without calcium and magnesium and dissociated using Accutase (130-091-221; STEMCEL Technologies, 4.

Vancouver, British Columbia, Canada). Cells were subsequently processed using autoMACS buffer (Milton Biotec, Bergisch Gladbach, Germany). Cells were fixed and infiltrated using a Cytofix/Cytoperm assay kit (554714; BD Biosciences) according to the manufacturer’s instructions. Cells were incubated overnight with a non-binding primary antibody in the dark at 4 °C, followed by incubation with fluorescence-labeled secondary antibodies for at least 3 h. Flow cytometry data were acquired and analyzed using BD FACSDiva software version 10.8 (BD Biosciences), and appropriate isotype matching was performed.

#### 5.2.7 Molecular Mechanism Study of the Drug-Loaded Microneedle Device in Long-Term Restenosis Intervention of Transplanted Veins

EdU and scratch assays were performed to evaluate the proliferation and migration of VSMCs. Western blotting and qPCR were used to determine the expression levels of glycolysis-related genes. The primer sequence is shown in Table S4.In addition, glucose uptake, lactate production, ATP generation, and ECAR of VSMCs were measured using the methods described above to evaluate the effects of the drug-loaded microneedle device on glycolytic activity. To further investigate the mechanisms associated with long-term restenosis, RNA sequencing, KEGG enrichment analyses, and heatmap generation were performed using samples derived from the surgical and control groups, as described above. Western blot analysis was subsequently performed to validate the protein expression levels of the selected genes in each group.

In addition, endothelial cell apoptosis assays were performed in each experimental group to determine the effects of the drug-loaded microneedle device on endothelial cell apoptosis.

#### 5.2.8 Statistical analysis

All data were analysed using the statistical analysis software SPSS 17.0.Data were presented as mean ± standard deviation. Comparison between two groups was conducted by Student’s t-test. Comparison among multiple groups was analysed by single-factor analysis of variance (ANOVA). Fisher’s least significant difference (LSD) test was also performed for two group comparisons. A value of P < 0.05 was considered statistically significant.Uniformly label the range of p-values in the figure (* p<0.05, * * p<0.01, * * p<0.001)

## Acknowledgements

The project was awarded the National Natural Science Foundation of China (82070364); Research and Cultivation Program of Beijing Municipal Hospital (PX2024026); Funded by the Beijing Municipal Natural Science Foundation of China and the Beijing Municipal Education Commission (KZ202010025044).

## Data Availability Statement

Data are available upon reasonable request

Received: ((will be filled in by the editorial staff))

Revised: ((will be filled in by the editorial staff))

Published online: ((will be filled in by the editorial staff))

## Supporting Information

Supporting Information is available from the Wiley Online Library or from the author.

---

**Figure S1.**
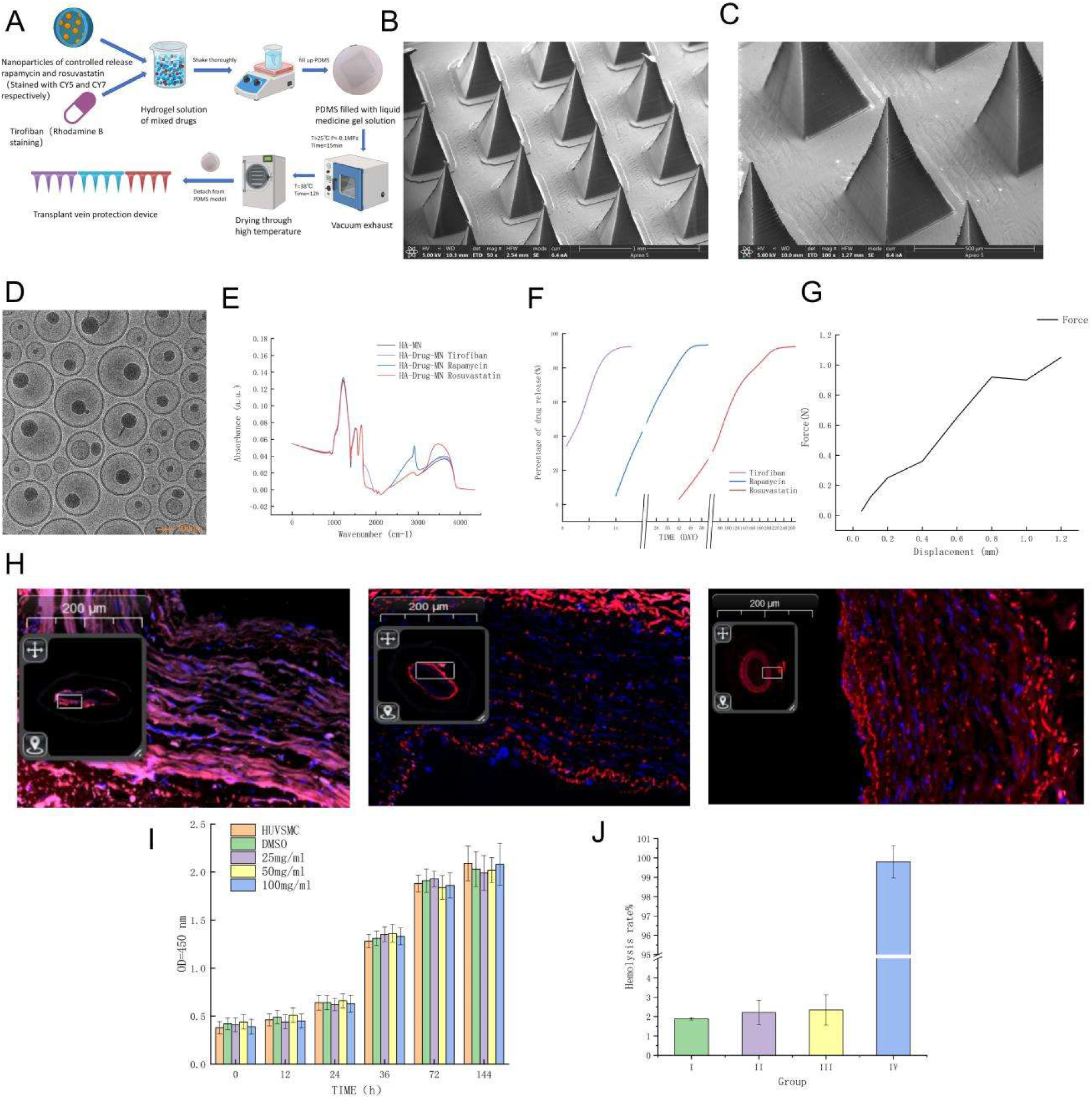
Preparation process of drug loaded microneedle device A, TEM results of drug loaded microneedle device C, TEM results of drug loaded nanoparticles D, infrared results of drug loaded microneedle device E, drug release curve of drug loaded microneedle device G, mechanical property curve of drug loaded microneedle H, fluorescence staining of drug loaded microneedle for treatment of human great saphenous vein

